# Enhanced Recognition of a Herbal Compound Epiberberine by a DNA Quadruplex-Duplex Structure

**DOI:** 10.1101/2024.05.14.594047

**Authors:** Xuan Zhan, Liping Deng, Yun Lian, Zhiyu Shu, Yunong Xu, Xinyi Mai, Manchugondanahalli S. Krishna, Chi Xiong, Rongguang Lu, Anni Wang, Shiyao Bai, Yingyi Xu, Jie Ni, J. Jeya Vandana, Zi Wang, Yuqing Li, Dongmei Sun, Shaohui Huang, Jingyan Liu, Gui-Juan Cheng, Song Wu, Ying-Chih Chiang, Goran Stjepanovic, Cheng Jiang, Yong Shao, Gang Chen

## Abstract

The small molecule epiberberine (EPI) is a natural alkaloid with versatile bioactivities against several diseases, including cancer and bacterial infection. EPI can induce the formation of a unique binding pocket at the 5′ side of a human telomeric G-quadruplex (HTG) sequence Q4, resulting in a nanomolar binding affinity (*K*_D_ approximately 26 nM) with significant fluorescence enhancement upon binding. It is important to understand (1) how EPI binding affects HTG structural stability and (2) how enhanced EPI binding may be achieved through the engineering of the DNA binding pocket. In this work, the EPI binding-induced HTG structure stabilization effect was probed by a peptide nucleic acid (PNA) invasion assay in combination with a series of biophysical techniques. We show that the PNA invasion-based method may be useful for the characterization of compounds binding to DNA (and RNA) structures in physiological conditions without the need to vary the solution temperature or buffer components, which are typically needed for structural stability characterization. Importantly, the combination of theoretical modeling and experimental quantification allows us to successfully engineer the Q4 derivative Q4-ds-A by a simple extension of a duplex structure to Q4 at the 5′ end. Q4-ds-A is a superb EPI binder with a *K*_D_ of 8 nM, with the binding enhancement achieved through the preformation of a binding pocket and a reduced dissociation rate. The tight binding of Q4 and Q4-ds-A with EPI allows us to develop a novel magnetic bead-based affinity purification system to effectively extract EPI from *Rhizoma coptidis* (Huang Lian) extracts.

---

Small molecule-nucleic acid complex formation is important for various biological processes and biotechnological applications. G-quadruplexes, consisting of guanine-rich DNA/RNA sequences, are stabilized by layers of Hoogsteen quadruples and metal ion coordination interactions^1-3^. Human telomeric DNA G-quadruplexes (HTGs), formed by repeating sequence of d(TTAGGG)_n_, may form multiple conformations depending on the sequence and the presence of monovalent cations (Na^+^, K^+^)^4-14^. The molecular recognition and targeting of HTG are of significant importance because HTG structure formation in human cells may be closely related to telomere lengthening and thus the impact on cancer evolution and aging^15-18^. In addition, recent efforts to discover aptamers selectively binding to small molecule fluorescent light-up probes often reveal that G-quadruplexes in combination with other structural motifs may provide an ideal structural scaffold for accommodating small molecules^19-27^.

An HTG sequence Q4, d(TTAGGG)_4_TTA (**Figures 1 and 2, Table 1**), is a fluorescent light-up sensor with a nanomolar binding affinity (∼26 nM) for a natural compound isoquinoline alkaloid epiberberine (EPI, **Figure 1A**)^28, 29^. The fluorescence enhancement may be due to a binding-induced reduction of intramolecular vibrational relaxation for the excited state (aggregation induced fluorescence emission (AIE) effect), as observed for an isomer of EPI, berberine (BER)^30-32^. The small molecule EPI shows biological activities in the inhibition of tumor growth^33^, urease activity^34^, and cytochrome P450 activity^35^ and in regulating serum cholesterol levels^36^. NMR structural studies revealed that EPI specifically recognizes the hybrid-2 type HTG formed in K^+^ solution^29^. Specifically, the previous NMR structural studies suggest that EPI binding to HTG Q4 can induce the formation of a hybrid-2 type conformation with a deep binding pocket at the 5′ side (see **Figure 1B**), through the formation of a quasi-triad plane with A3 and intercalation between the top G-tetrad plane and the triad plane formed by T2, T13, and A15 (**Figure 1B**, see the sequence numbering in **Figure 2**). The EPI-HTG Q4 complex formed in the Na^+^ solution is significantly weakened due to the absence of the binding pocket with the Na^+^-specific HTG conformation^28, 29^.

**Table 1.**
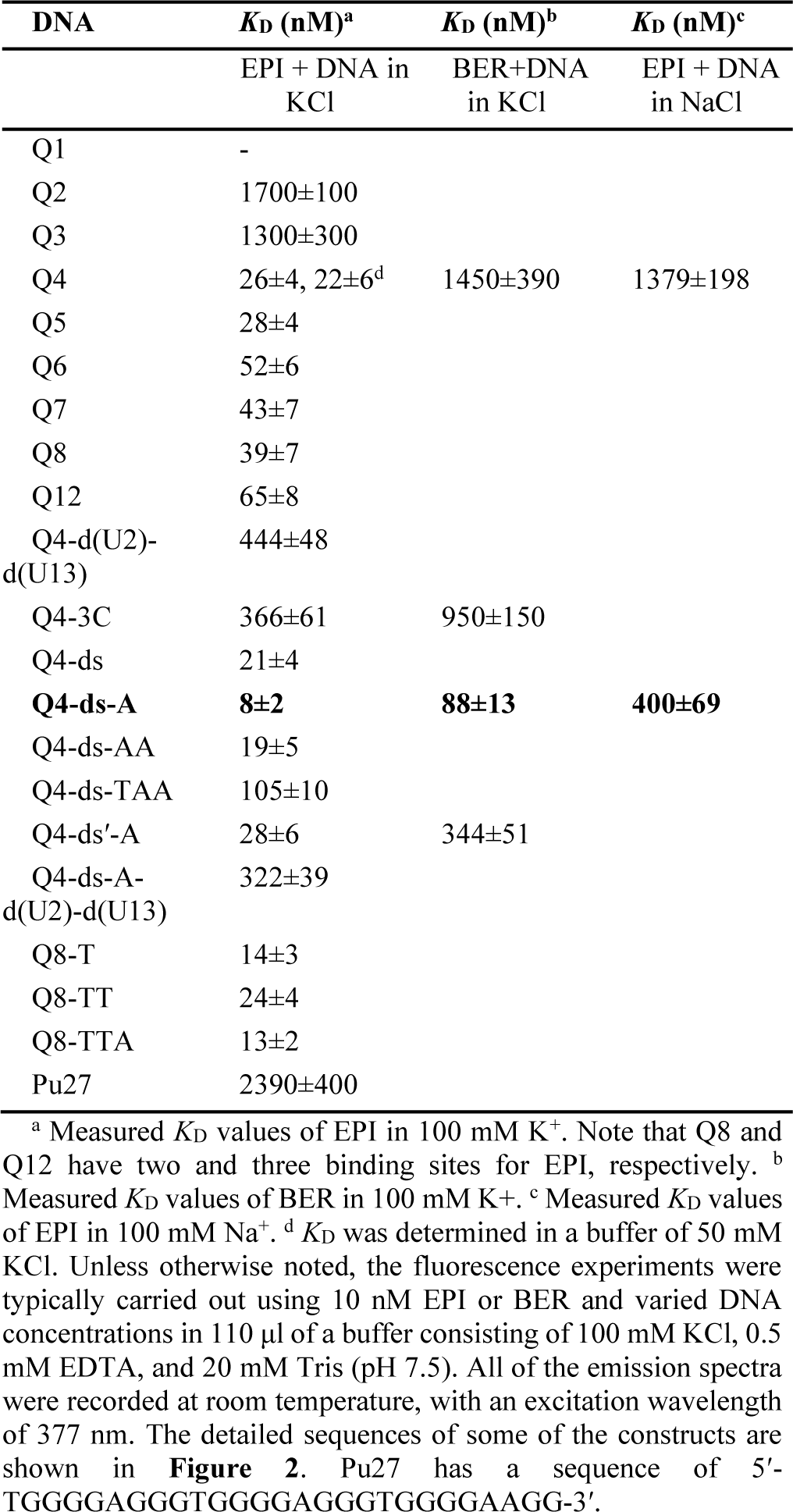
Measured *K*_D_ values by steady-state fluorescence titration assay for DNAs Qn: (TTAGGG)_n_TTA (n = 1-8, 12) and other DNAs.

**Figure 1.**
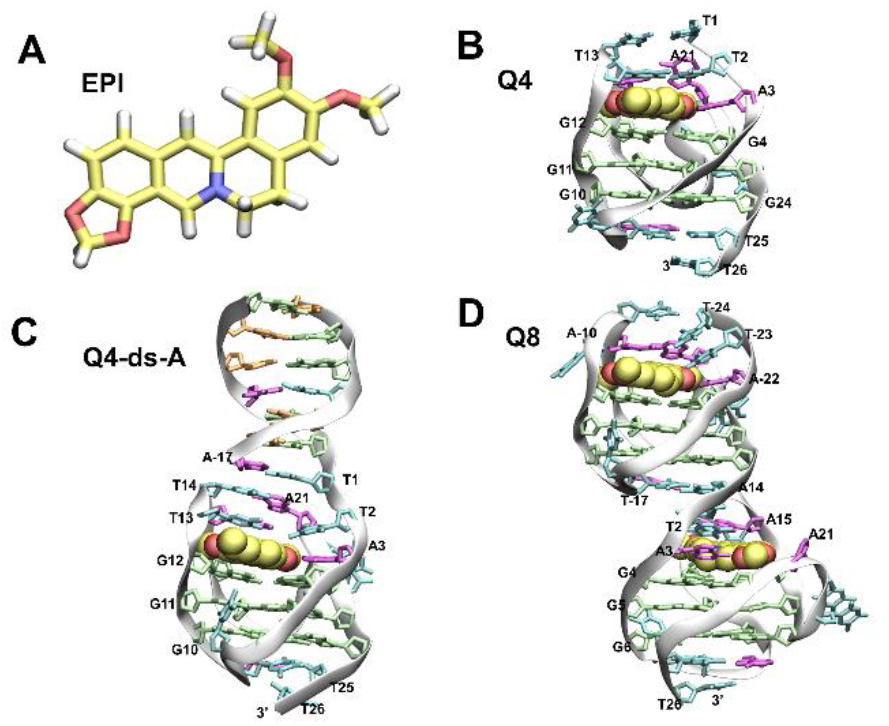
Structures of EPI and DNA-EPI complexes. **(A)** Chemical structure of epiberberine. The carbon, oxygen, nitrogen, and hydrogen atoms and corresponding chemical bonds are shown in yellow, red, blue, and white, respectively. (**B-D**) NMR structure and modelled structures of DNA-EPI complexes. The nucleotides G, A, T, and C are depicted in green, magenta, cyan, and orange, respectively. In the NMR and modeled structures, the 3′ terminal A residue of the DNAs is missing compared to the DNAs listed in **Figure 2. (B)** NMR structure of the Q4-EPI complex (PDB ID: 6ccw)^29^. Q4 forms a hybrid-2 type conformation with a deep binding pocket for EPI. **(C)** Modeled structure of Q4-ds-A in complex with EPI. **(D)** Modeled structure of Q8 in complex with EPI. The second binding pocket of EPI was used for free energy calculation. For the latter two panels, there are no available experimental structures. For panels C and D, depicted are the structures of the major cluster centroids, obtained from three runs of 100 ns MD simulations.

**Figure 2.**
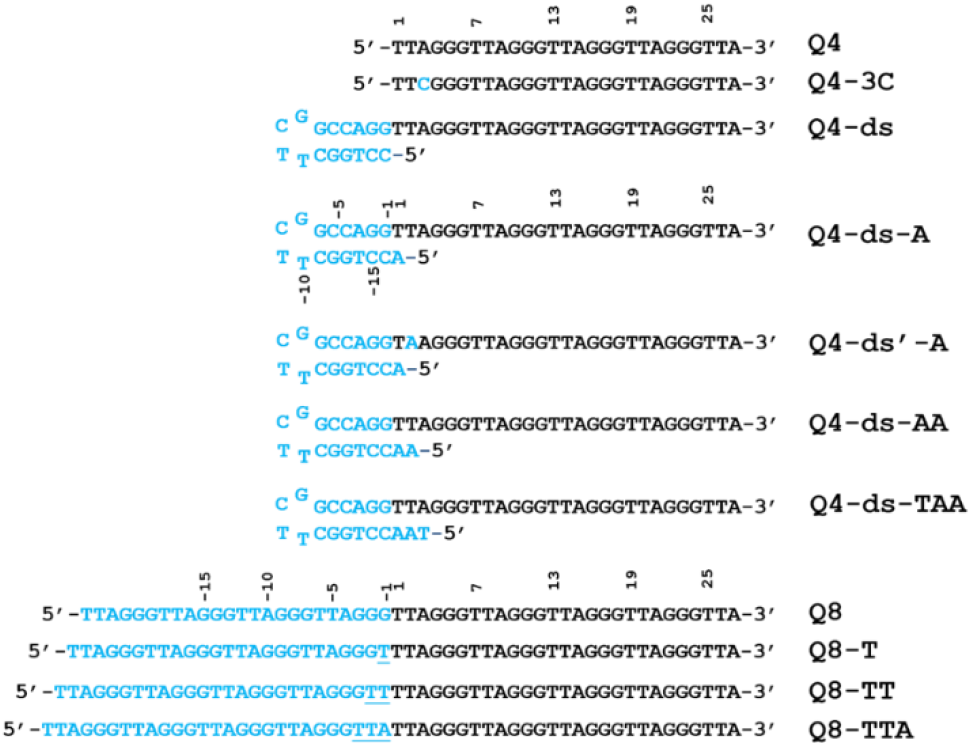
DNA sequences constructed based on Q4. The sequences shown in cyan are engineered based on the Q4 sequence; the engineered residues are labelled with negative numbers for representative sequences.

Quantifying the stabilization effects of nucleic acid structures by small molecule binding is critical for understanding nucleic acid functions and regulations by small molecule ligands. Typically a thermal unfolding process is needed to probe the stabilization of biomolecular structures upon small molecule binding, which, however, may not reflect the real situation in the cell. Antisense peptide nucleic acids (asPNAs) are useful for binding to complementary nucleic acid sequences through the formation of PNA-DNA or PNA-RNA duplexes^37, 38^. We hypothesize that competitive binding by asPNA and small molecules to HTG may allow for the characterization of HTG structure stabilization by small molecule binding. In this work, we experimentally verified, by an array of quantitative methods, including fluorescence, nondenaturing polyacrylamide gel electrophoresis, and native mass spectrometry, the competitive binding between a series of potential HTG-binding small molecules and asPNA. We show that the asPNA invasion method may be employed to quantify how small molecule binding stabilizes HTG structures.

Furthermore, designing and discovering nucleic acid sequences for enhanced recognition of small molecules may pave the way for the development of advanced nucleic acid-based techniques for applications in biosensing and other biotechnologies. In this work, engineering of Q4 aided by computer simulations results in a novel DNA construct Q4-ds-A, which contains an extension of a duplex structure at the 5′ end of Q4. Q4-ds-A shows improved binding of EPI and enhanced colocalization with EPI in the cell. Finally, we demonstrate that Q4-ds-A is useful for affinity purification of EPI from semi-purified plant products (*Rhizoma coptidis* extract).

## EXPERIMENTAL SECTION

### Materials and Reagents

Experimental reagents and solvents were used without further purification. DNA oligomers were purchased from Suzhou Biosyntech. The PNA oligomers were synthesized by a standard solid-phase peptide synthesis method with the monomers purchased from ASM.

### Nondenaturing Polyacrylamide Gel Electrophoresis

The nondenaturing polyacrylamide gel electrophoresis (PAGE, 20% (acr:bis=29:1)) experiments were performed using an incubation buffer A: 25 mM Tris-HCl, pH 7.5, 100 mM NaCl, 1 mM EDTA or incubation buffer B: 25 mM Tris-HCl, pH 7.5, 100 mM KCl, 1 mM EDTA, in the presence or absence of 1 or 2 µM EPI. PNA and DNA oligomers with and without small molecules were dried in Speed-Vac at 30°C and dissolved in an incubation buffer. The solutions were heated at 95°C for 5 min followed by slow cooling (∼2 h). Then, the samples were incubated at 4°C for 4 h or overnight. The running buffer A contained 1× TBE and 100 mM NaCl, pH 7.5 (150 V, 50 mA, 6 h). The running buffer B contained 1× TBE and 100 mM KCl, pH 7.5 (150 V, 50 mA, 6 h). The final concentration of Q4 was 2 µM (25 nM for Cy5-Q4). The gels for the samples without Cy5 label or EPI were stained in 350 mL of 1 µg/mL ethidium bromide (EB) solution for 30 minutes and scanned in a Typhoon instrument at an excitation wavelength of 532 nm.

### Magnetic Bead-Based Native Purification

The magnetic beads were prepared in-house. Briefly, spherical magnetic particles with carboxyl groups (with a diameter of 270 nm) were synthesized by a hydrothermal method^39^. Specifically, 2.7 g FeCl_3_ · 6H_2_O, Na acrylate, and polyethylene glycol were dissolved in 80 ml ethylene glycol under magnetic stirring. The obtained solution was transferred to a Teflon-lined stainless-steel autoclave and sealed to heat at 200 °C for 8 h. The autoclave was then cooled to room temperature and washed several times with water and anhydrous ethanol to clarify the supernatant. Fe_3_O_4_ magnetic beads were harvested with aid of magnetic rack, followed by drying at 60 °C. Magnetic beads with epoxy group modification on the surface were obtained by dispersing 10 mg of magnetic beads into absolute ethanol, with the addition of 3 μL (3-glycidyloxypropyl)trimethoxysilane (GPTMS) under stirring at room temperature for 3 h. The products (expoy funcitonalzied magnetic beads) were subsequently cleaned with ethanol and water^40^. The high density of epoxy groups on the surface of magnetic beads allows mild and efficient covalent coupling with thiolated nucleic acids. The shaking processes were performed on a multipurpose shaker (QB210, Kylin-Bell Lab Instrument). The supernatant after magnet application was further analyzed by EPI fluorescence emission and HPLC. For the HPLC experiments, an Agilent 1260 HPLC and a C18 column were used, with buffer A composed of water with 0.1% TFA and buffer B containing HPLC-grade acetonitrile. The gradient was set as follows: 0-2 min: 90% A + 10% B; 2-6 min: 80% A + 20% B; 6-10 min: 70% A + 30% B; 10-14 min: 60% A + 40% B; 14-18 min: 50% A + 50% B; 18-20 min: 5% A + 95% B; 20.1 min: 90% A + 10% B; 20.1-22 min: 90% A + 10% B. The column temperature was set at 45 °C.

## RESULTS AND DISCUSSION

### EPI binding to natural human telomeric DNA sequences

EPI shows a “light-up” behavior upon tight and specific binding to HTG sequences, e.g., Q4 (**Figure 3**) ^28, 29^. In this work, we further explored how varying the natural human telomeric sequence and length may affect binding with EPI (**Table 1**). The steady-state fluorescence titration data (obtained by titrating DNA into EPI, see **Figures 3 and S1**) reveal that, in a KCl solution, Q4 exhibits the strongest binding, with Q1 showing no observable binding: Q4 (26 nM) < Q5 (28 nM) < Q7 (43 nM) < Q8 (39 nM) < Q6 (52 nM) < Q12 (65 nM) < Q3 (1.3 μM) < Q2 (1.7 μM) (the numbers after the letter Q indicate the n values in d(TTAGGG)_n_TTA, **Table 1**).

**Figure 3.**
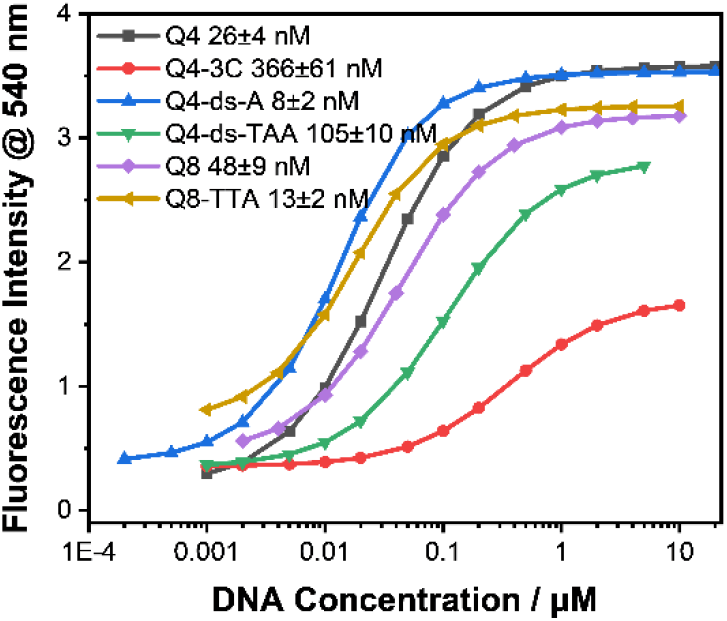
*K*_D_ measurement of EPI by steady-state fluorescence titration. All of the fluorescence experiments were carried out using 10 nM EPI and varied DNA concentrations in 110 μl of a buffer consisting of 100 mM KCl, 0.5 mM EDTA, and 20 mM Tris (pH 7.5). All of the emission spectra were recorded at room temperature, at an excitation wavelength of 377 nm. For DNAs Q8 and Q8-TTA, we used the concentration of HTG (twice the concentration of Q8 and Q8-TTA) to determine the *K*_D_ values.

We further characterized the complex formation by native mass spectrometry (MS) assay^41-43^. The native MS data suggest that the EPI-Q4 complex is highly stable with the major binding ratio of EPI and Q4 of 1:1, in NH_4_^+^ solution (a mimic of K^+^), with a small fraction showing 1:2 binding (**Figure S2**). It can’t be excluded that the two complexes with 1:1 and 1:2 binding ratios may have different ionization properties. We used NH_4_OAc instead of KCl for the native MS study, as NH_4_ ^+^ is a mimic of K^+^ and is suitable for the MS study. Consistently, the UV absorbance-detected thermal melting data suggest that EPI is an effective stabilizer of G-quadruplexes (Q4-Q8, Q12) in K^+^ solution with an increase in the melting temperature (Δ*T*_m_) of 11-15 °C (**Figure S3**).

We then tested whether EPI can be used to stain HTG structures in a nondenaturing PAGE experiment. Indeed, compared to ethidium bromide (EB) prestaining, HTG sequences (e.g., Q4-Q12) are prestained more efficiently by EPI, with both incubation and running buffers containing KCl (**Figure S4**). However, changing to NaCl-containing incubation and running buffers causes no staining of the gel bands except for Q8 and Q12. The PAGE experiment data further confirm that EPI fluorescence light-up effect is due to tight and specific binding to K^+^ cation-specific hybrid-2 type conformation of HTG. Interestingly, Q8 migrates faster than Q6 and Q7, suggesting that two quadruplex units form within Q8, with a more compact structure compared to Q6 and Q7. The data show that EPI may be useful for imaging and probing complex HTG structure formation with varied telomeric repeat numbers.

We further probed EPI-Q4 binding in the cellular environment. Our confocal microscopic imaging data revealed that Q4 labelled with a 3′-end Cy5 (**Figure S5**) is indeed colocalized with EPI (**Figure S6A-D**). The red, green, and blue fluorescence signals correspond to Cy5-labelled Q4, EPI, and Hoechst, respectively. The orange signals in the merged image result from the co-localization of EPI and Cy5-lablled Q4. Thus, the intracellular imaging data indicate that EPI indeed binds to Q4 tightly, which implies that Q4 forms the hybrid-2 type structure in the cell. Furthermore, the cell-free telomerase activity assay data show that EPI may inhibit telomere extension catalyzed by telomerase (**Figure S7**), possibly through stabilizing the hybrid-2 type G-quadruplex structure potentially formed by the newly synthesized telomeric repeat sequence^44-46^.

### PNA invasion assay for the quantification of the quadruplex structure stabilization effect of EPI

asPNAs show strong binding to complementary DNAs and RNAs and have been used for probing tertiary structure formation in RNA pseudoknots^47^ and RNA structural rearrangement^48^. The asPNAs containing the complementary sequence of human telomeric sequence have been used to successfully invade HTG structures^49-52^. We carried out asPNA invasion (**Figure 4A, Figure S8**) assay to quantitatively probe the binding and stabilization of HTG by a series of small molecules including EPI.

**Figure 4.**
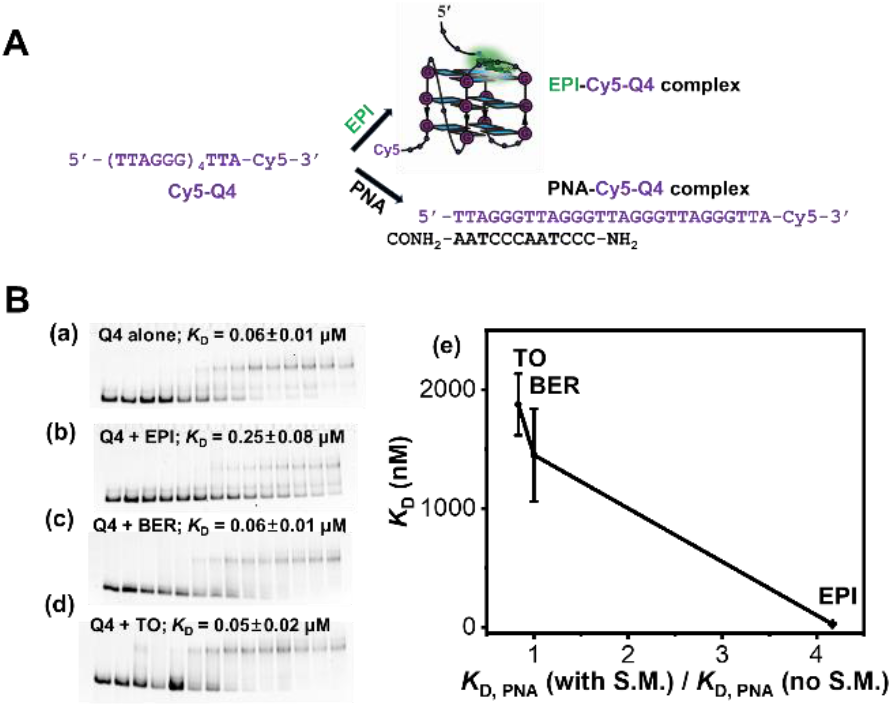
PNA invasion assay. (**A)** Schematic of the competitive binding of EPI and PNA to Cy5-labelled Q4 (Cy5-Q4). **(B)** Nondenaturing PAGE study of PNA (NH_2_-Lys-CCCTAACCCTAA-CONH_2_) titrating to Cy5-labelled Q4 (10 nM) in the absence (a) and presence of 1 µM EPI (b), 1 µM BER (berberine, an isomer of epiberberine, c) 1 µM and thiazole orange (TO, d). The incubation buffer contained 25 mM Tris-HCl (pH 7.5), 100 mM KCl, and 1 mM EDTA. PNA concentrations are 0-20 µM. (e) The relationship between small molecule binding with Q4 (*K*_D_, Y-axis) and the ratio of PNA invasion *K*_D,PNA_ with and without small molecules determined by PAGE assay (X-axis).

We employed native MS to investigate the binding stoichiometry of the three-component system composed of Q4, EPI, and asPNA. The complexes were assembled in NH_4_^+^ solution (a mimic of K^+^ suitable for the MS study) and directly ionized using an offline nanospray source. We used a 7-mer asPNA (NH_2_-Lys-ACCCTAA-CONH_2_) in the native MS study (**Figure S2**). In a mixture of 10 nM Q4 and 1 μM asPNA, we observed MS peaks for the complexes of 1:1 Q4-asPNA and 1:2 Q4-asPNA. Interestingly, no Q4-asPNA complex with a higher ratio was observed under this experimental condition. In a mixture of 10 nM Q4, 1 μM EPI and 1 μM asPNA, we observed MS peaks for the complexes of 1:1 Q4-EPI, 1:1 Q4-asPNA, and 1:2 Q4-asPNA but no peak for the ternary complex of Q4, EPI, and asPNA was detected. Taken together, these results support the competitive binding of EPI and asPNA to Q4 (**Figure 4A**).

We employed the PNA invasion method for probing the binding and stabilization of Q4 structure by EPI and other potential small molecule ligands using a nondenaturing PAGE assay (**Figure 4B**). As there was no significant gel shift resulting from the invasion of Q4 by the 7-mer PNA (NH_2_-Lys-ACCCTAA-CONH_2_) in the PAGE assay (**Figures S5 and S8**), we decided to make a 12-mer PNA (NH_2_-Lys-CCCTAACCCTAA-CONH_2_) as the Q4 invader. We measured the binding affinities of the 12-mer PNA invading Q4 construct covalently labelled with Cy5 dye at its 3′ end (**Figures S5 and S8**), with and without the presence of small molecules. Consistently, the *K*_D,PNA_ value for PNA invasion increased significantly, from 0.06 ± 0.01 µM to 0.25 ± 0.08 µM upon the addition of 1 µM EPI (**Figure 4B**). The ratio of *K*_D,PNA_ (with EPI) vs *K*_D,PNA_ (without EPI) is 4.2 (**Figure 4B**). Compared to EPI, berberine (BER) and thiazole orange (TO) show relatively weakened binding to Q4 as measured by steady-state fluorescence titrations (**Figures S9 and S10**). Remarkably, the addition of 1 µM each of BER or TO results in no observable change in the apparent *K*_D,PNA_ value of PNA invasion, with the ratios of *K*_D,PNA_ (with small molecule) vs *K*_D,PNA_ (without small molecule) being approximately 1.0 (**Figure 4B**). Our proof-of-principle data show that asPNA competitive binding through Q4 structure invasion is a convenient method for probing tight binders of Q4.

We next applied fluorescence correlation spectroscopy (FCS) to further probe the competitive binding of EPI to a preformed HTG-asPNA complex in NaCl and KCl buffers, respectively (**Figure S11**). FCS measures the diffusion coefficients (D) of fluorescently labeled molecules, and the D-values can then be used to calculate the corresponding hydrodynamic radii of the molecules^53^. In this FCS study, we again used the Q4 construct covalently labelled with Cy5 dye at its 3′ end (**Figure S5**), which forms the (Cy5-Q4)-asPNA duplex in sample solutions containing saturating amounts of unlabeled asPNA. Then, various concentrations of EPI were added into the preformed (Cy5-Q4)-asPNA duplex; the competitive binding of EPI to Cy5-Q4 thus results in a gradual replacement of the (Cy5-Q4)-asPNA duplex with (Cy5-Q4)-EPI complex (**Figure 4A**). The FCS titration data indicate that the apparent hydrodynamic radius of the former molecular complex is slightly larger than that of the latter with the corresponding diffusion coefficients measured to be 137.2 and 142.2 μm^2^s^−1^, respectively. Based on the FCS data, we derived the apparent dissociation constants (*K*_D_) of 47 nM and 172 nM for EPI binding to Cy5-Q4 in the KCl and NaCl buffers, respectively (**Figure S11**). The data are consistent with the binding affinities determined by the steady-state fluorescence titration experiments, with EPI having a tighter binding affinity to Q4 in KCl buffer (26 nM) than in NaCl buffer (1467 nM) (**Table 1**).

### Probing the EPI binding pocket of Q4 through bimolecular duplex formation at the 5′ side

With multiple telomeric repeats, HTG structure formation may be dynamic, with the G-quadruplex structure formed at 5′ side, 3′ side, or in the middle. To avoid such complexity, we made a Q5-5′ss sequence with non-telomeric sequences added at the 5′ side of Q5 (**Figure 5**). We designed a series of antisense DNAs with varied lengths complementary to the 5′ side sequence to probe the importance of the 5′ end single strand region for EPI binding (**Figure 5**).

**Figure 5.**
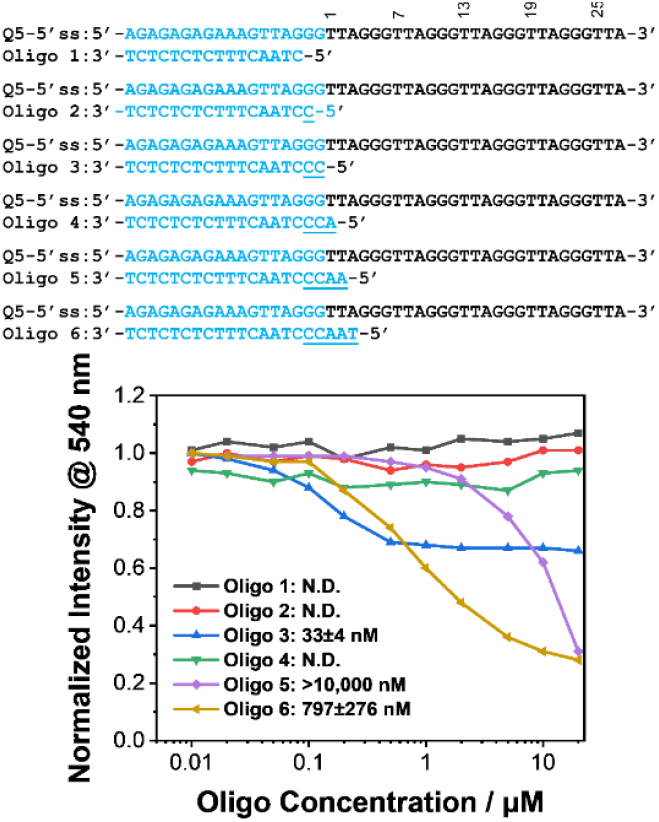
Characterization of the 5′ binding pocket by forming a bimolecular 5′ duplex structure. A series of oligonucleotides (oligos 1-6) were titrated into a preformed complex of EPI (0.2 μM) and Q5-5′ss (0.2 μM). All of the EPI fluorescence experiments were carried out using varied oligo concentrations (0-20 μM) in a buffer consisting of 25 mM Tris-HCl, pH 7.5, 100 mM KCl, and 1 mM EDTA (pH 7.5). All of the emission spectra were recorded at room temperature, at an excitation wavelength of 377 nm. N.D.: Binding affinity for the oligonucleotide was not determined as no significant change in EPI fluorescence emission intensity was observed.

As expected, the addition of DNA oligomers 5 and 6 results in a decrease in EPI fluorescence, presumably due to the duplex formation at the 5′ side resulting in the disruption of the EPI binding pocket as revealed previously^29^. A small decrease in EPI fluorescence intensity was observed upon the addition of oligomer 3, indicating that the hybridization of oligomer 3 (resulting in the formation of a structure similar to Q4-ds, **Figure 2**) may slightly affect the binding of EPI. Interestingly, titration of oligomer 4 into the preformed Q5-5′ss-EPI complex (resulting in the formation of a structure similar to Q4-ds-A, **Figure 2**) caused no response in EPI fluorescence intensity. The data on bimolecular DNA constructs suggest that the formation of the 5′ side duplex to Q4 with a two-nucleotide “single-stranded” linker sequence of 5′TA3′ may not significantly affect the local structural environment of EPI.

### Engineering Q4 sequences for enhanced binding towards EPI

We next constructed unimolecular DNA constructs to further probe the EPI binding pocket of Q4. Since the binding pocket of EPI is located at the 5′ end of Q4, we focused on engineering the sequences/structures at/near the 5′ end. The sequence modifications (**Figure 2** and **Table 1**) were mainly focused on: **i)** The sequence extension at the 5′ end with double-stranded DNA stem, which resulted in DNA constructs Q4-ds, Q4-ds-A, and Q4-ds-TAA. **ii)** Replacing d(A3), d(T2) and d(T13) in Q4 with d(C3), d(U2)/d(A2) and d(U13), which generated the constructs Q4-3C, Q4-d(U2)-d(U13), Q4-ds′-A, and Q4-ds-A-d(U2)-d(U13). **iii)** Inserting T, TT, and TTA between the fourth and fifth repeats of TTAGGG formed in Q8 to generate derivatives Q8-T, Q8-TT, and Q8-TTA.

We evaluated the binding of the newly constructed sequences by the steady-state fluorescence titration method (**Table 1, Figure 3, Figures S12-S28**). A single A3C mutation (Q4-3C) results in a significant weakening of the binding (from 26 nM to 366 nM). The result suggests that C is too small compared to A in forming an ideal hydrogen bond with EPI and stacking interaction within the DNA as shown by the previous NMR study^29^. Q4-d(U2)-d(U13) also shows a significantly weakened binding affinity (444 nM), suggesting the importance of the triad formed by A15, T2 and T13.

Remarkably, the single-chain construct Q4-ds-A shows the strongest binding (*K*_D_ = 8 nM) among the tested DNA sequences (**Figures 2** and **3** and **Table 1**). Interestingly, a single-stranded spacer of at least one residue (A3) is needed for tight binding (**Figure 2** and **Table 1**), which is consistent with the previous binding and NMR data^28, 29^, suggesting the importance of A3 in forming a hydrogen bond with EPI.

Interestingly, intracellular confocal imaging data (**Figure S6**) suggest that Q4-ds-A shows better colocalization with EPI, than Q4 and the control G-quadruplex sequence Pu27 (**Table 1**)^54, 55^.

We characterized the EPI binding kinetics using bio-layer interferometry (BLI), through the immobilization of 3′ end biotin-labelled DNA constructs on surface. The BLI data (**Figure S29**) show that Q4 has a *K*_D_ of 384 nM, which is higher than that by steady-state fluorescence measurement (26 nM, **Table 1**). In consideration of the differences in the experimental setup (with DNA immobilized on surface versus free in solution), the *K*_D_ of 384 nM value obtained through kinetic measurement is reasonable. The association rate of 135,000 M^−1^S^−1^ is relatively slow for a small molecule, probably due to the fact that a conformational rearrangement of Q4 is needed for EPI binding. We observed that Q4-3C has a significantly weakened binding (*K*_D_ = 17,600 nM) with reduced *k*_on_ and increased *k*_off_. Interestingly, Q4-ds-A exhibits a strengthened binding (*K*_D_ = 185 nM) with slightly increased *k*_on_ and moderately decreased *k*_off_.

We further asked whether extending the Q4 at the 5′ end with another quadruplex with a suitable spacer may improve the binding affinity. In the original Q8 sequence (**Figures 1 and 2)**, the middle binding pocket may have two competing conformations, as the TTA sequence is present at the 3′ side of the first Q4 and 5′ side of the second Q4. Indeed, we observed improved binding upon inserting single-stranded spacers of T, TT, and TTA sequences into Q8 (**Table 1**). Our data suggest that the natural repeating sequences of human DNA telomere are not ideally evolved for stable formation of multiple hybrid-2 type HTG in K^+^, and engineered telomere sequences/structures may allow improved G-quadruplex formation and ligand binding.

### Theoretical modelling

We carried out Molecular Dynamics (MD) simulations of the EPI-DNA complexes to gain insights into the varied binding strengths of EPI with the DNAs. We calculated the binding free energies of Q4, Q4-ds, Q4-ds-A, Q4-3C, and Q8 to EPI using umbrella sampling. The calculated free energies and measured binding affinities show a correlation (*R*^2^ = 0.67, **Figure 6A**). To probe the molecular determinants for the binding of EPI to various DNA constructs, we carried out a detailed analysis of the initial structures of the complexes and the trajectories from the umbrella simulations. We observed that the binding pocket (local structures above the G-quadruplex) in Q4-EPI complex (**Figure 6B**) became disordered soon after the EPI was pulled out of the binding pocket, suggesting that the binding pocket only forms in the presence of EPI, which agrees well with the previous NMR study.^29^ In the Q4-3C construct with an A3C mutation (**Figure 6C**), a hydrogen bond can still form between C3 and EPI, similar to the hydrogen bond formed between A3 and EPI in the Q4-EPI complex. The reduced stability of (Q4-3C)-EPI complex (**Figure 6A**) may be due to the relatively small size of C compared to A, resulting in reduced stacking interactions.

**Figure 6.**
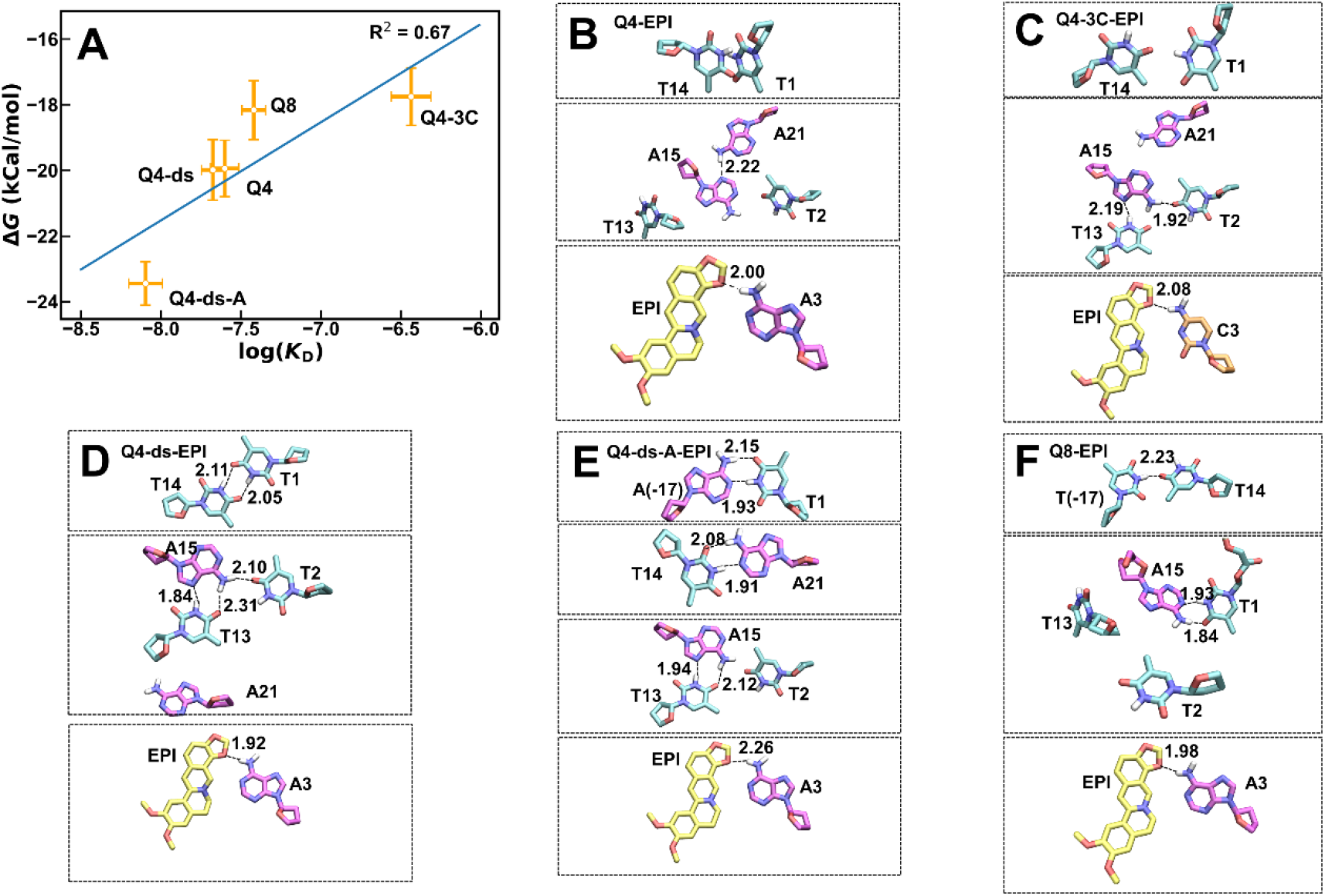
Comparison of the calculated binding free energy of EPI and simulated structures of the binding pockets of EPI. **(A)** Comparison between calculated binding free energies and experimentally measured log(*K*_D_). Errors owing to the finite sampling were calculated and depicted in the figure. Panels **B-F** show the detailed interactions of the binding pockets of **(B)** Q4-EPI; **(C)** Q4-3C-EPI; **(D)** Q4-ds-A-EPI; **(E)** Q4-ds-EPI, and **(F)** Q8-EPI. The first binding pocket of Q8 for EPI is essentially the same as that of Q4 and thus is not shown. The calculated free energy for Q8 is an average of the two binding pockets.

As expected, Q4-ds and Q4 (**Figure 6A,B,D**) show similar calculated binding free energy, with the binding pocket showing flexibility. The MD simulation data suggest that the hybrid duplex-quadruplex structure Q4-ds-A may involve structural rearrangement for accommodating EPI (**Figures 1 and 6E**). The binding pocket in Q4-ds-A (**Figure 6E**), which is different from that of Q4 (**Figure 6B**), is significantly more stable. The local structure above the G-quadruple surrounding EPI was formed even after EPI was pulled away. The enhanced local structural stability could be attributed to the T2·T13·A15 triple and T14·A21 base pair that lie above the binding pocket (**Figure 6E**). Although a T2·T13·A15 triple or T2·A15·A21 triple was widely observed in simulation trajectories for all engineered Q4s (**Figure 6B-F**), the newly formed T14·A21 pair stacked above the triple was only observed in Q4-ds-A. It is probable that the newly formed T14·A21 base pair stacked right below the T1-A(−17) Watson-Crick pair (see **Figure 2** for the structural scheme) greatly improves the binding of Q4-ds-A by stabilizing the binding pocket. Without the extra A(−17) residue (as present in Q4-ds-A), T1 is dynamic and may pair with or be stacked with T14 as observed for other constructs. Q8 has two EPI binding pockets, with the first binding pocket of Q8 for EPI essentially the same as that of Q4. We observed the second binding pocket is dynamic, although the hydrogen bond between A3 and EPI is still stably formed (**Figure 6F**). Taken together, our experimental and computational modelling data suggest that fixing T1 by base pairing with A(−17) as observed for Q4-ds-A seems to be essential to achieve an improved binding with EPI.

### Extraction of Epiberberine from *Rhizoma coptidis* extracts using Q4 and Q4-ds-A

*Rhizoma coptidis* is a Chinese medicinal herb with potential biomedical applications for treating a wide range of diseases^56^. The effective alkaloids consist of BER as the major component and other compounds such as EPI, BER, palmatine (PAL), columbamine (COL), and coptisine (COP) (**Figure 7A**)^56^. The extraction and separation of these alkaloids are of significant importance for their versatile applications in pharmacological applications. We reason that the tight binding between EPI and natural Q4 and engineered DNAs may be utilized for affinity purification of EPI from *Rhizoma coptidis*, which may be complementary to the existing HPLC method^57^. Leveraging the differential binding of EPI with HTG-containing structures in K^+^ and Na^+^ solutions (**Table 1**), we designed a magnetic bead (MB)-based simple approach (**Figure 7B**) to extract EPI from *Rhizoma coptidis* extract.

**Figure 7.**
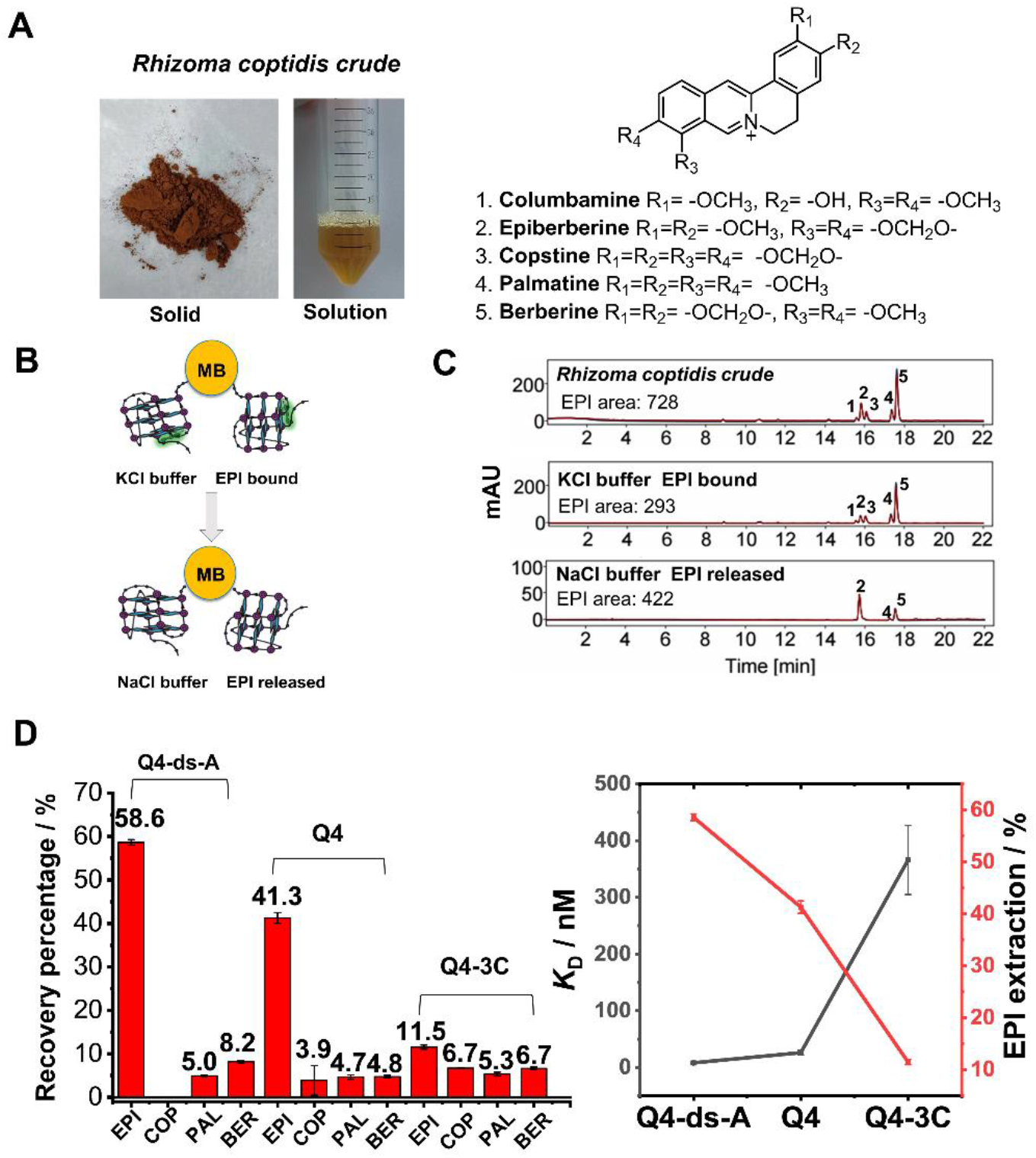
Application of the engineered DNA for affinity purification of EPI. **(A)** Pictures of studied *Rhizoma coptidis* crude samples (solid and solution state (1 mg/mL)) and chemical structures of five main alkaloid small molecules found in sample crude and HPLC traces of COL, EPI, COP, PAL and BER correspond to peaks 1-5, displayed in panel C. **(B)** General principle for the affinity purification of EPI via a simple magnetic beads assay. EPI is in the bound state in the KCl environment and is released in the NaCl environment, due to the formation of two different quadruplex conformations, resulting in significantly different binding affinities with EPI (**Table 1**). **(C)** HPLC traces of the supernatants (from top to down) in *Rhizoma coptidis* crude sample (0.05 mg/mL, 10 μL HPLC injection, top), treatment with KCl buffer and shaking for 2 hours (middle), and treatment with NaCl buffer and shaking for 10 h (bottom). More operational details can be seen in the experimental section and supporting information section. A summary of all HPLC traces is shown in **Table S1. (D)** Affinity purification recovery rates of the alkaloids based on HPLC analyses. (Left panel) Comparison of three different DNAs for the magnetic bead-based affinity purification of the *Rhizoma coptidis* crude sample. (Right panel) The *K*_D_ value (DNA-EPI binding) inversely correlates with the recovery percentage.

In the MB-based affinity extraction/purification approach, we covalently immobilized DNAs (with a thiol group functionalized at the 3′ end, **Figure S8**), on epoxylated MB. Using thiolated Q4 and pure EPI, we optimized the protocol for DNA covalent immobilization, EPI binding in KCl, impurity washing in KCl, and EPI elution in NaCl (**Scheme S1, Figures S30-S35**). For a pure EPI solution, the optimized affinity purification protocol shows a net EPI recovery yield of 85% (**Figure S33**) with an EPI binding solution of 50 mM KCl, followed by impurity washing in 50 mM KCl, and EPI elution by 5 min of heating at 95°C followed by incubation in 200 mM NaCl. In a solution with 50 mM KCl, we observed that under UV light, free Q4-ds-A can effectively light up EPI in solution (**Figure S36**). However, with Q4-ds-A covalently immobilized on the MB, no fluorescence of EPI was observed. Clearly, EPI binds specifically to Q4-ds-A structure on MB.

We next applied the MB-based affinity purification strategy to purify EPI from *Rhizoma coptidis* extract, which contains a mixture of COL, EPI, COP, PAL, and BER as revealed by HPLC (**Figure 7A,C**). We compared the extraction efficacies among Q4, Q4-ds-A, and Q4-3C, with their *K*_D_ values for EPI (25 nM, 8 nM, and 366 nM) and BER (1379 nM, 88 nM, and 950 nM) (**Table 1**). Q4-ds-A showed the best EPI recovery (58.6%), followed by Q4 (41.3%) and Q4-3C (11.5%), which is consistent with their affinities with EPI (**Figure 7C,D, Figure S37, Table S1**). We further used diluted HCl (pH 3) instead of 200 mM NaCl for the final elution step using Q4-ds-A as the capture probe. We observed an EPI recovery of 62.4% (**Table S1**). It is important to note that COL, EPI, and COP are clustered in the HPLC, which may cause difficulty in purifying EPI. However, no observable amount of COL or COP was detected after affinity purification using the engineered DNA Q4-ds-A (**Figure 7C,D**). A small amount of recovery of PAL and BER (**Figure 7C,D**) may be avoided by further optimization of the affinity purification protocol. Our affinity purification approach may be complementary to purification solely based on HPLC. The nucleic acid-based affinity purification approach may be further optimized and applied to the extraction of many other valuable natural products.

## CONCLUSIONS

In this work, we developed a PNA invasion-based assay to quantify the stabilization effect of nucleic acid structures upon binding by small molecule ligands. Using the HTG structure as a model system, we reveal that the binding affinity of small molecules to HTG correlates with the asPNA invasion of the small molecule-HTG complexes. There is great potential in employing the PNA-invasion method for screening small molecule drug candidates targeting DNA and RNA structures^58^. Natural nucleic acid structures provide great molecular scaffolds for further molecular engineering for improved binding properties. Significantly, our molecular engineering efforts aided by MD simulations result in Q4-ds-A, which shows a superb binding affinity towards EPI with a *K*_D_ of 8 nM. The work provides new insights into the fundamental principles governing molecular recognition of EPI by DNA. We further developed a magnetic bead-based affinity purification approach using the engineered DNA Q4-ds-A. Such affinity purification allows the purification of hard-to-purify components in natural products solely based on HPLC. The advantage of the native affinity purification method is that no harmful organic solvents are needed. It is conceivable that with the development of rational engineering and selection (e.g., aptamer) approaches, one may develop multiplexed systems for the affinity purification and detection of multiple compounds.

## ASSOCIATED CONTENT

### Supporting Information

This material is available free of charge via the Internet at http://pubs.acs.org. Experiments of steady-state fluorescence study, thermal Melting, confocal microscopy imaging assay, native mass spectrometry, telomerase activity assay, FCS measurement, BLI measurement, DNA structure modeling, MD simulation, umbrella sampling for free energy calculation are well described in the supporting information.

### Notes

A patent application has been filed based on the work described in this article.

## ACKNOWLEDGMENT

The work was supported by The Chinese University of Hong Kong, Shenzhen (CUHK-Shenzhen) University Development Fund (to G.C., C.J., G.S., and Y.-C.C.), Guangdong Provincial Basic and Applied Basic Research Fund Project-Youth Funding (2022A1515110577 to X.Z.), Ganghong Young Scholar Development Fund (PhD Studentship) (to L.D.), National Natural Science Foundation of China General Program (22177098 to G.C.), fund from Shenzhen-Hong Kong Cooperation Zone for Technology and Innovation (HZQB-KCZYB-2020056, to G.C.), the Leading Talent Program of Science and Technology Innovation in Zhejiang (Grant No. 2020R52022, to Y.S.), the National Natural Science Foundation of China (to G.S., 31950410540) and Foreign Youth Talent Program from State Administration of Foreign Experts Affairs (to G.S., QN2021032004L), 2022 Joint Fund of School of Medicine and The Second Affiliated Hospital of The Chinese University of Hong Kong, Shenzhen (CUHK-Shenzhen) (to J.L. and G.C.), and Shenzhen Science, Technology and Innovation Committee for the Shenzhen Key Laboratory Scheme (ZDSYS20220507161600001). The authors thank the advanced mass spectrometry facility (KMS) of Kobilka Institute of Innovative Drug Discovery, the Chinese University of Hong Kong, Shenzhen for the support.

