## Supplement Material for "Enhanced Recognition of a Herbal Compound Epiberberine by a DNA Quadruplex-Duplex Structure"

| Entry | Content | Pages |
| --- | --- | --- |
| **Materials and Method** | Materials and experimental details | 3-6 |
| **Figure S1** | Fluorescence binding study of EPI binding to natural telomere sequences | 7 |
| **Figure S2** | Native Mass Spectrometry data. | 8 |
| **Figure S3** | Thermal melting curves of natural telomere sequences | 9 |
| **Figure S4** | Comparison of DNA structure staining by EPI and ethidium bromide (EB) | 9 |
| **Figure S5** | Labelled DNAs and PNAs used | 10 |
| **Figure S6** | Intracellular colocalization assay of EPI with DNA | 11 |
| **Figure S7** | TRAP assay data of telomerase activity | 12 |
| **Figure S8** | PAGE assays of a 7-mer PNA binding to DNA | 12 |
| **Figures S9-S28** | Fluorescence binding data for $K_D$ value determination | 12-21 |
| **Figure S29** | BLI measurement | 22 |
| **Scheme S1** | Optimization experiments conducted for EPI extraction | 23 |
| **Figures S30-S35** | Optimization data for EPI extraction | 23-27 |
| **Figure S36** | Visualization of fluorescence light-up effect of EPI | 28 |
| **Figure S37** | Procedure for alkaloid extraction | 28 |
| **Table S1** | HPLC characterization of EPI extraction protocol | 29 |
| **References** | References for the Supporting Information | 30 |

### MATERIALS AND METHODS

#### Materials

Experimental reagents (Aldrich) and solvents were used without further purification. DNA oligos were purchased from Suzhou Biosyntech. The PNA monomers were purchased from ASM.

#### Steady-State Fluorescence Study

The fluorescence emission spectra were recorded using a 1 cm square quartz cuvette on a Varian Cary Eclipse fluorescence spectrophotometer. The fluorescence experiments were typically carried out using 10 nM EPI and varied DNA concentrations in a buffer solution consisting of 100 mM NaCl/KCl, 0.5 mM EDTA, and 20 mM HEPES (pH 7.5), unless otherwise noted. The samples were prepared by slow cooling from 95°C to room temperature, followed by the addition of G-quadruplexes. The solutions were then vortexed for 10 min, left at room temperature for 1 h, and incubated at 4°C overnight. The excitation wavelength was 377 nm. All of the emission spectra were recorded from 450 nm to 700 nm at room temperature.

#### Thermal Melting

The thermal melting experiments were conducted with the use of an eight-microcell cuvette using the Shimadzu UV-2550 ultraviolet-visible spectrophotometer. The absorbance values at 295 nm with the temperature increasing from 15 to 95°C followed by the temperature decreasing from 95 to 15°C were recorded. The temperature ramp rate was 0.2°C/min. G-quadruplexes were formed by annealing in an incubation buffer (25 mM HEPES pH 7.5, 100 mM KCl, 1 mM EDTA). The concentration of DNA was 2 μM. Then, 10 μM EPI was added after annealing. Data were normalized at low temperature, and the melting temperatures were determined based on the Gaussian fits of the first derivatives of the curves.

#### Nondenaturing Polyacrylamide Gel Electrophoresis

The nondenaturing polyacrylamide gel electrophoresis (PAGE, 20% PAGE (acr:bis=29:1)) experiments were performed using an incubation buffer A: 25 mM Tris-HCl pH 7.5, 100 mM NaCl, 1 mM EDTA or incubation buffer B: 25 mM Tris-HCl pH 7.5, 100 mM KCl, 1 mM EDTA. 1 or 2 μM EPI. G-quadruplex Q4 (DNA oligos: 5′-TTAGGGTTAGGGTTAGGGTTAGGGTTA-3′) with or without Cy5 labelled and PNA were dried in Speed-Vac at 30°C and dissolved in the incubation buffer. Running buffer A contained 1× TBE and 100 mM NaCl, pH=7.5 (150 V, 50 mA, 6 h). Running buffer B contained 1× TBE and 100 mM KCl, pH=7.5 (150 V, 50 mA, 6 h). The final concentration of Q4 was 2 μM (25 nM for Cy5-Q4), and the final concentrations of PNA/DNA oligos were 0 – 50 μM. The gels for the samples without EPI were stained in 350 mL of 1 μg/mL ethidium bromide (EB) solution for 30 minutes and scanned in a Typhoon instrument at an excitation wavelength of 532 nm. For the annealing of the samples, PNA and RNA samples were dried, followed by the addition of buffer (25 mM Tris-HCl, 100 mM KCl/NaCl, 1 mMEDTA, pH=7.5), 95°C for 5 min and slow cooling (~2 h). Then, the samples were incubated at 4°C for 4 h or overnight.

#### Confocal Microscopy Imaging Assay

HEK293T cells were seeded on 14 mm circle microscope cover glasses in 24-well plates at a density of $5\times10^4$ per well and grown in high glucose DMEM medium at 37°C for 24 h. Then, the cells were treated with EPI at different concentrations alone for 12 h. The medium was then removed and the cover glasses were washed with PBS 3 times. The cover glasses were then placed on slides sealed with 50% glycerol for observation under a confocal microscope. The apparatus was a Zeiss LSM900 confocal microscope. The excitation wavelengths for detecting EPI, Hoechst and Cy5 were 488 nm, 405 nm and 640 nm, respectively.

**Native Mass Spectrometry**

For direct infusion with the nano source, the Offline Nano ES kit (Thermo Scientific) was used. Samples were loaded in a Borosilicate Offline Emitter (Thermo Scientific), injected by means of the static air pressure device and measured for 5 min with a full MS scan on a Q Exactive HFX Hybrid Quadrupole-Orbitrap mass spectrometer (Thermo Scientific). For ionization in negative mode, a spray voltage of 0.8 kV and capillary temperature of 350°C were used. For all measurements, the AGC target value was $3e^6$, the maximum injection time was set to 200 ms, and the resolution setting was 60,000 and 10 µscans. The source-induced collisional dissociation (SID) energy was set to 50 eV, supporting desolvation and declustering and greatly improving spectral quality. The source-CID energy was carefully tuned and kept below values leading to sample dissociation. In parallel, all data were acquired and verified with SID turned off. Data analysis and spectra deconvolution were performed by Thermo Scientific FreeStyle 1.8 SP1 software.

**Telomerase Activity Assay**

Telomerase activity was determined by TRAP assay (telomeric repeat amplification protocol) using a TRAPeze XL telomerase detection kit (Millipore, USA) per the manufacturer's instructions. Telomerase was extracted from HEK293-T cells using CHAPS buffer containing RNase inhibitor at 1:100. The telomerase template was first elongated with cell extracts containing telomerase and different concentrations of EPI at 30°C for 30 min, and then amplified by PCR (94°C-30 s, 59°C-30 s, 72°C-60 s for 36 cycles. Then, 72°C-3 min, 55°C-25 min, 4°C for infinite hold). The samples were then loaded on native PAGE gels (10%, Acr:Bis = 19:1) and run at 400 V in 1× TBE buffer for 3 h at 4°C. The gel was stained by SYBR Gold (Sangon Biotech, China) at 1:10000 for 15 min and then washed in $H_2O$ for 15 min. The image was taken using Typhoon at 532 nm excitation wavelength. The 56 bp band in every lane is the internal control. The intensity of the band ladder (starting from 61 bp) and the internal control was determined using ImageJ software.

**Fluorescence Correlation Spectroscopy**

FCS experiments were carried out using a CorTector SX100 benchtop fluorescence correlation spectrometer produced by LightEdge Technologies Limited (Zhongshan, Guangdong, China). The instrument is equipped with an Olympus 60X NA 1.2 water immersion objective and two continuous wave lasers (488 nm and 638 nm) from Pavilion Integration Corporation (Suzhou, Jiangsu, China). To perform an FCS experiment, ~50 µl of sample solution was added to a high-precision coverslip (CG15CH; Thorlabs, Newton, New Jersey, USA) that was balanced directly on top of the objective lens. The instrument was first calibrated using a 10 nM Atto655 carboxylic acid (ATTO-TEC GmbH, Siegen, Germany) solution, which allows the optimization of the confocal optical pathway of the instrument and determines the FCS volume (<1 fL) using the dye's reported diffusion coefficient of 426 $\mu m^2 s^{-1}$ in 25°C water [1]. The (Cy5-Q4)-asPNA complex was formed by mixing 1 nM Cy5-Q4 and 100 nM asPNA, followed by incubation in a 37°C water bath for 2 h. Increasing amounts of EPI samples were added to the (Cy5-Q4)-asPNA solution. Competitive EPI binding to Cy5-Q4 then causes the gradual replacement of (Cy5-Q4)-asPNA with (Cy5-Q4)-EPI. The former molecular complex has a smaller diffusion coefficient (hence a larger hydrodynamic radius) compared to that of the latter complex, which can be distinguished by FCS measurements. Thus, competitive binding curves of EPI to the (Cy5-Q4)-asPNA complex were constructed, from which the apparent dissociation constant ($K_D$) was derived [2a].

Briefly, the auto-correlation decay curves were fitted using the following equation:

$$G(\tau)=\frac{1}{N}\cdot\frac{1-T+Te^{-\frac{\tau}{\tau_T}}}{1-T}\cdot\left(F_P\cdot\frac{1}{\left(1+\frac{\tau}{\tau_P}\right)\cdot\sqrt{1+\frac{\tau}{\tau_P S^2}}}+F_{\mathrm{E}}\cdot\frac{1}{\left(1+\frac{\tau}{\tau_{\mathrm{E}}}\right)*\sqrt{1+\frac{\tau}{\tau_{\mathrm{E}} S^2}}}\right)\quad Eq. 1$$

where N is the average number of the Cy5 labeled G4 molecules in the FCS volume element; T and $\tau_T$ are the fraction of Cy5 fluorophore in the triplet state and its characteristic relaxation time for the dye's single-triplet dynamics respectively; $F_P$ and $\tau_T$ are the fraction of Cy5-Q4 molecules in the Cy5-Q4-asPNA binding state and the molecular complex's characteristic diffusion correlation time respectively; $F_E$ and $\tau_E$ are the fraction of Cy5-Q4 molecules in the Cy5-Q4-EPI binding state and the molecular complex's characteristic diffusion correlation time respectively; S = $z_0/r_0$ is the structure parameter of the FCS volume element, with $z_0$ and $r_0$ being the axial and radial dimensions of the volume element respectively. The T and $\tau_T$ values were independently determined using a 10 nM Cy5 dye solution, and were subsequently fixed in Eq. 1 for subsequent analysis of sample data. $F_P$ and $F_E$ are related to the average number of the Cy5-Q4-asPNA and Cy5-Q4-EPI molecule complexes in the FCS volume element (*i.e.*, $N_P$ and $N_E$) using the following equation:

$$\frac{N_P}{N_E}=\left(\frac{\lambda_E}{\lambda_P}\right)^2*\frac{F_P}{F_{\mathrm{E}}}\quad Eq. 2$$

$\lambda_P$ and $\lambda_E$ are the fluorescence brightness of the Cy5-Q4-asPNA and Cy5-Q4-EPI molecular complexes respectively, which can be experimentally determined using samples with zero or saturation concentrations of EPI respectively. The competitive binding curve is defined by:

$$F_b=\frac{N_E}{N_P+N_E}=\frac{1}{\frac{N_P}{N_E}+1}\quad Eq. 3$$

The competitive binding curves were fitted using the following equation to obtain the apparent dissociation constant for Cy5-Q4 and EPI binding:

$$F_b=\frac{p\cdot[EPI]}{K_D+[EPI]}+B\quad Eq. 4$$

where p and B are proportional constant and baseline respectively.

The Atto655 and Cy5 dyes were excited using a 638 nm laser, and the resulting fluorescence signals were selected by a T647lpxr dichromic mirror and an ET655lp long-pass filter before being detected by a single-photon counting avalanche photodiode (SPCM-AQRH-14; Excelitas Technologies, Waltham, Massachusetts, USA). Each FCS measurement lasted 10 seconds and was repeated 20 times. FCS data acquisition and data analysis were accomplished using the *Correlation Acquisition* and *Correlation Analysis* software developed by LightEdge Technologies.

**Bio-layer interferometry (BLI) characterization of binding kinetics of EPI**

Streptavidin SAXT probes (Gator) were used to determine the binding kinetics and affinities for EPI with various DNA constructs. The experiments and data analyses were carried out using GatorPrime system (Gator). The 3' biotin labeled Q4, Q4-3C and Q4-ds-A (Biosyntech) DNA samples were diluted by binding buffer (containing 100 mM KCl, 1 mM EDTA, 25 mM Tris-HCl, pH 7.5) into a final concentration of 200 nM. The concentrations of EPI are 10000, 5000, 2500, 1250, 625 and 0 nM. The association and dissociation time periods are both 20 seconds at 30℃.

**DNA Structure Modeling**

The Q4 structure in complex with EPI employed in the molecular dynamics (MD) simulations was taken from the Protein Data Bank (PDB code 6CCW)[3]. For Q4-ds, Q4-ds-A, Q4-3C and Q8, no experimentally resolved structures are available and thus were modeled based on the Q4 template. The former two models were built by linking the Q4-EPI complex to the associated B-DNA structures, generated by 3D-DART[4]The Q4-3C model was obtained by a direct mutation of d(A3) into d(C3), while the Q8 model was constructed by connecting two Q4 structures together. Within all the G-quadruplex structures, two potassium ions were placed in between the three G-quadruples.

**MD Simulation**

The MD simulations were performed using the GROMACS 2021 [5] package with the OL15 [6] parameters for DNA and the AM1-BCC charge model with GAFF2 [7] parameters for the ligand. The DNA-EPI complex was solvated in a cubic water box filled with TIP3P water molecules. The system was then further neutralized by potassium ions. The distance from the complex to the box edge was chosen to be at least 1.5 nm. The short-range cutoff for the nonbonded interaction was set to 1 nm, while the long-range electrostatic interaction was calculated by the particle-mesh Ewald (PME) method [8] with a spline order of 4 and a grid spacing of 0.12 nm. The temperature was controlled at 300 K and the pressure was controlled at 1 bar by the Langevin thermostat and Parrinello-Rahman pressure coupling, respectively. The integration time step was chosen to be 2 fs. Prior to the production run, all systems were first subjected to 50000-step energy minimization, 500 ps NVT equilibration and 500 ps NPT equilibration. Finally, 3 copies of 100 ns equilibrium run were performed. The DNA-EPI complex trajectories were clustered using GROMOS clustering algorithm [9] with an RMSD cutoff of 2.5 Å for Q4 and Q4-3C and 5 Å for others larger complex.

**Umbrella Sampling for Free Energy Calculation**

The initial structures used in umbrella sampling were taken from the major cluster centroid obtained in the MD simulations, except for Q4, where the Q4-EPI NMR structure is available. The umbrella sampling MD simulations were performed using the same parameters as described above, with a box of size 9 × 11 × 20 $nm^3$. First, EPI and G-quadraplex (top layer) were pulled apart using the pull code[10] with a pulling rate of 0.02 nm/ps, and the DNA was restrained. Then, 90 windows were used to sample along the reaction coordinate, i.e., the distance between center-of-masses of the two groups. Within each window, this distance was restrained by a harmonic potential with a force constant of 1000 kJ/mol/$nm^2$, and 30 ns of MD simulation with three layer of G-quadraplexes restrained was then performed for data collection. Finally, the potential of mean force (PMF) was calculated using the weighted histogram analysis method (WHAM)[11].

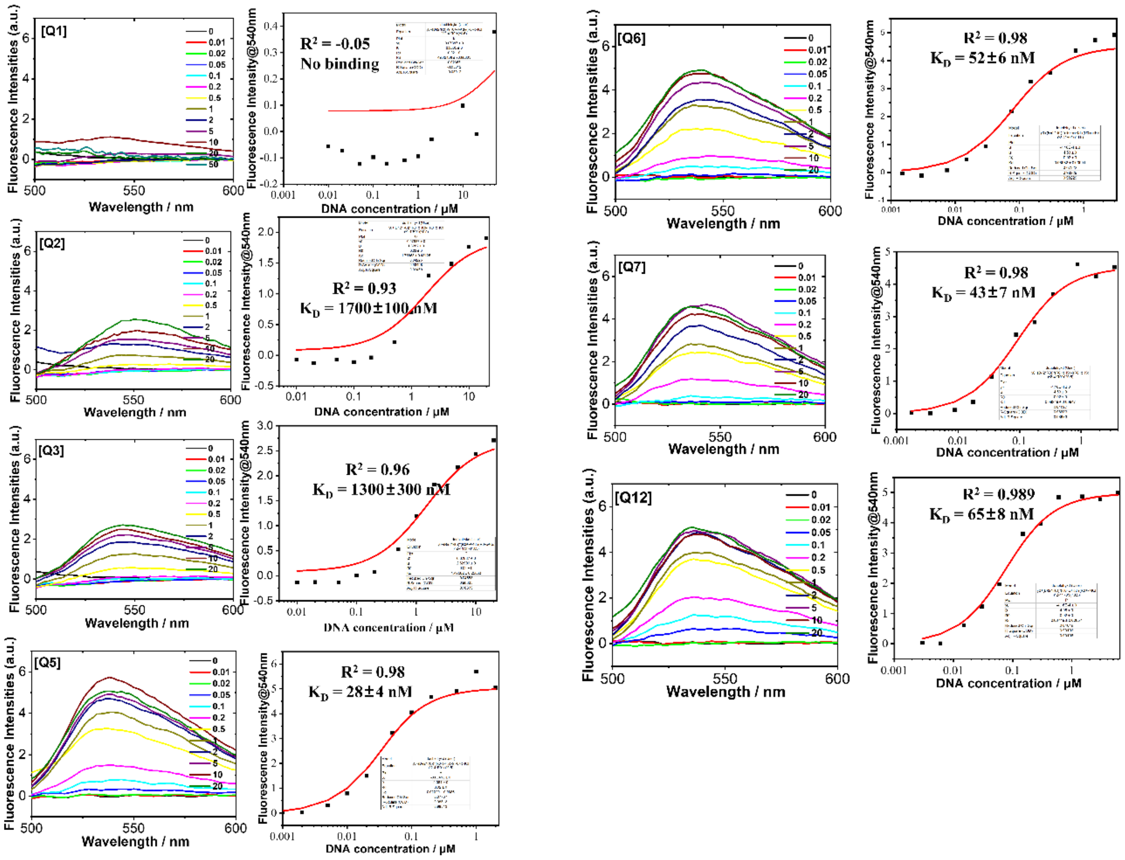


**Figure S1.** Fluorescence binding study of natural telomere sequences. Titration of Q1, Q2, Q3, Q5, Q6, Q7, Q8 and Q12 into EPI monitored by EPI fluorescence emission The concentration of EPI was 20 nM, the incubation buffer was: 25 mM Tris-HCl pH 7.5, 1 mM EDTA, 100 mM KCl. The $K_D$ fitting curves of the binding data are also shown. For Q1, no fitting was done.

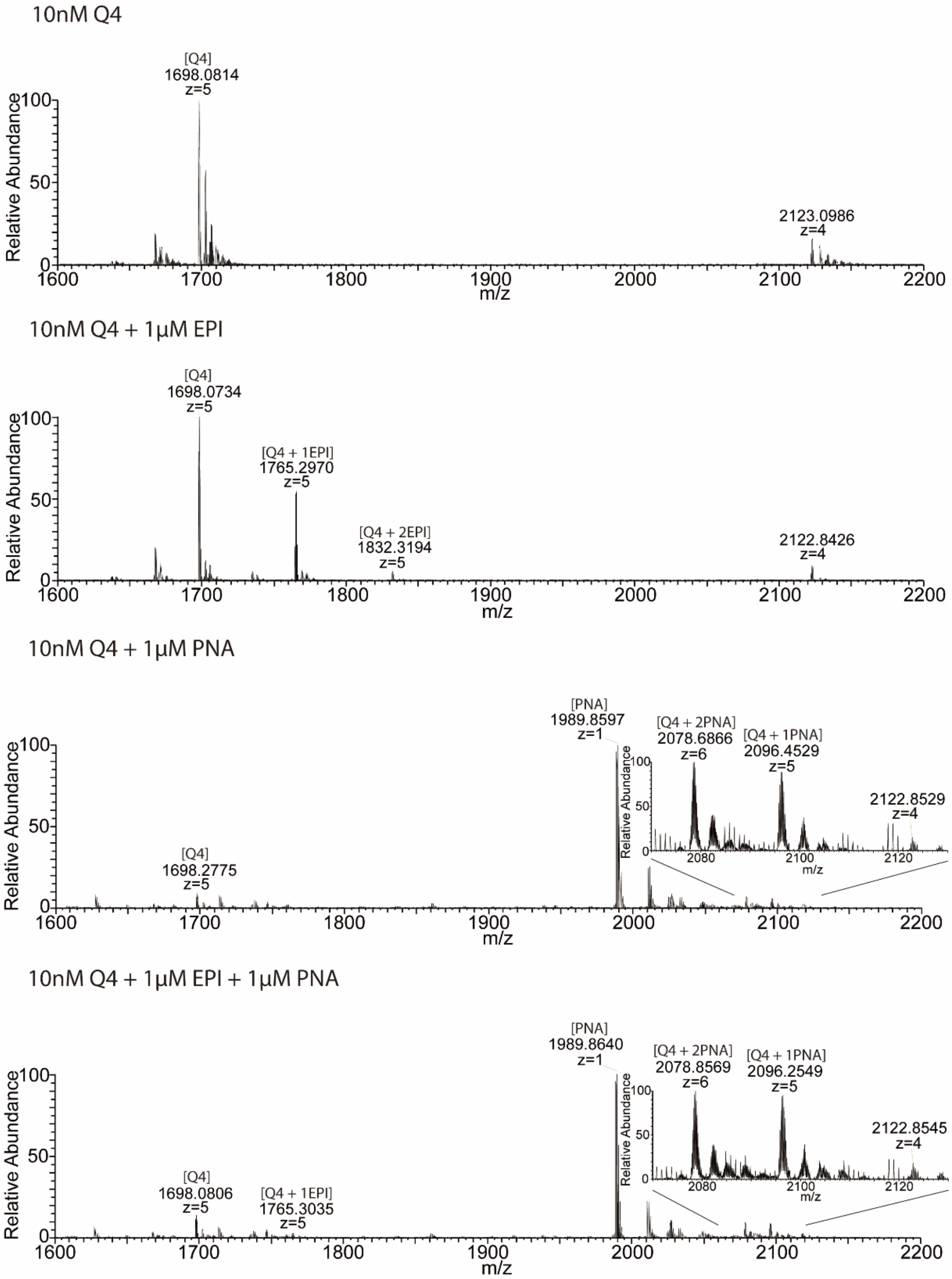


**Figure S2.** Native Mass Spectrometry. The native ESI mass spectra were obtained in $NH_4OAc$ (100 mM). The PNA used was $NH_2$-Lys-ACCCTAA-$CONH_2$.

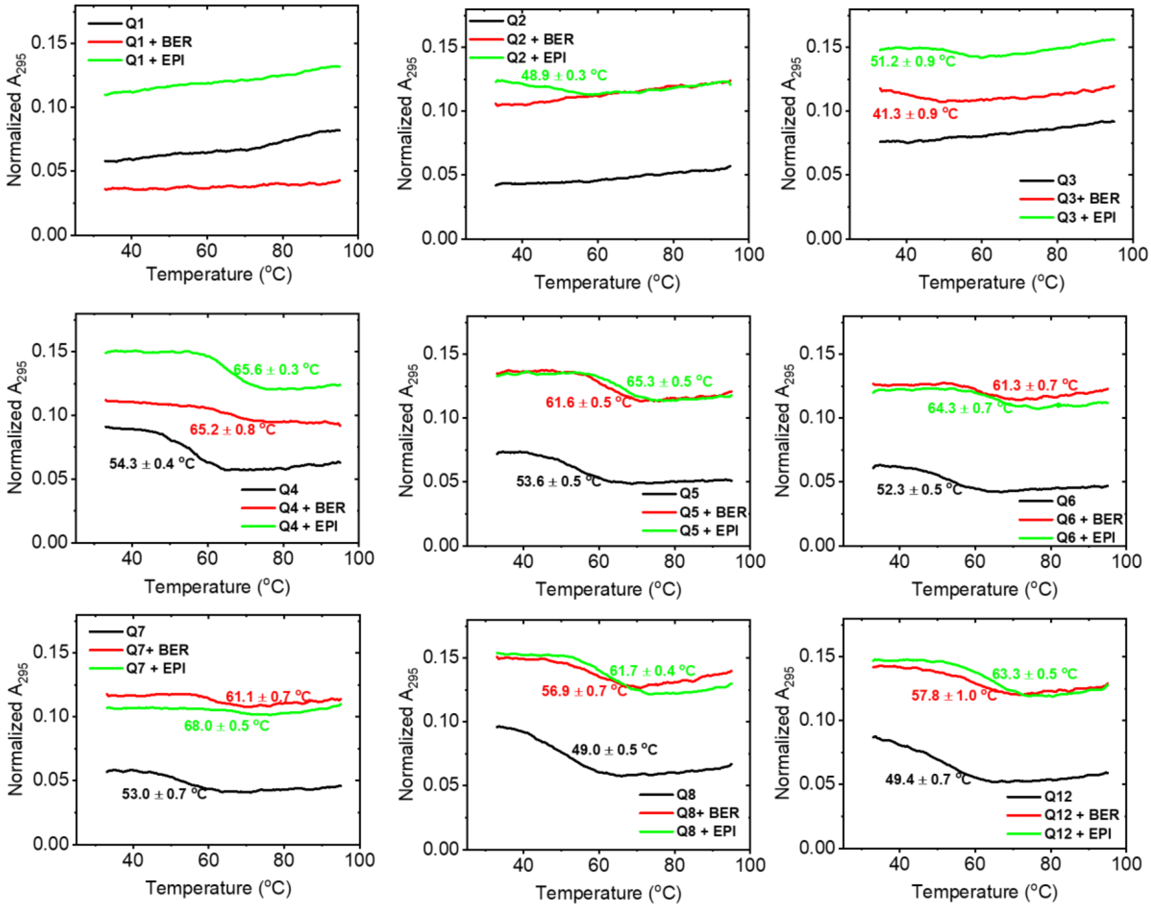

**Figure S3.** Thermal melting curves of natural telomere sequences in the absence and presence of EPI/BER. The incubation buffer: 25 mM HEPES pH 7.5, 100 mM KCl, 1 mM EDTA. The concentration of EPI was 10 µM. The concentration of DNAs was 2 µM.

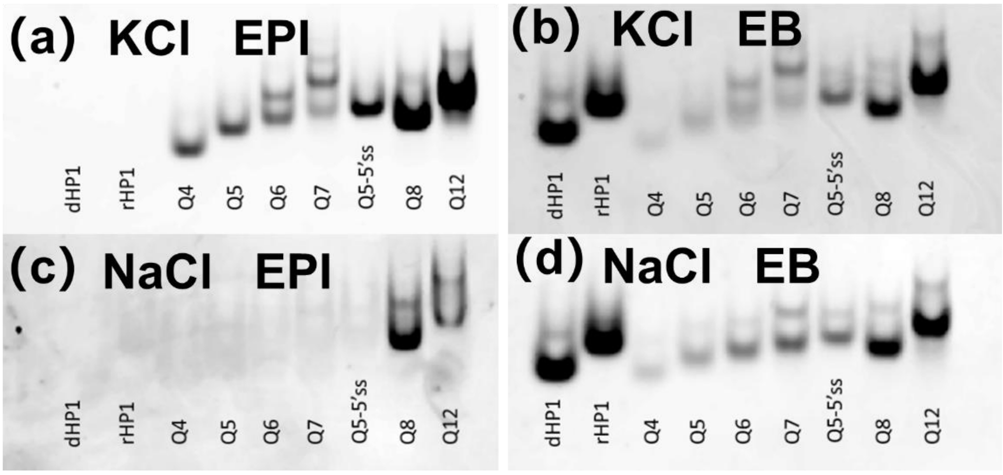


**Figure S4.** Comparison of DNA structure staining by EPI and ethidium bromide (EB). The incubation buffer was: 25 mM Tris-HCl pH 7.5, 100 mM KCl or 100 mM NaCl, 1 mM EDTA. The Q5-5′ss sequence is 5′-AGAGAGAGAAAG(TTAGGG)$_4$TTA-3′ (**Figure 5**).

(a)

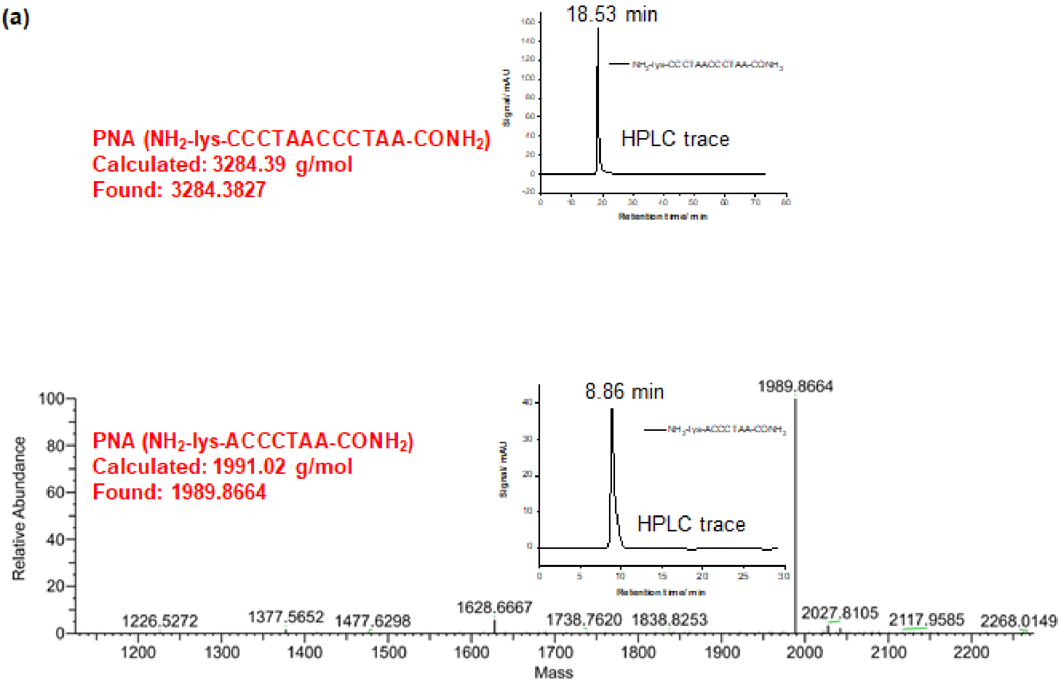


(b)

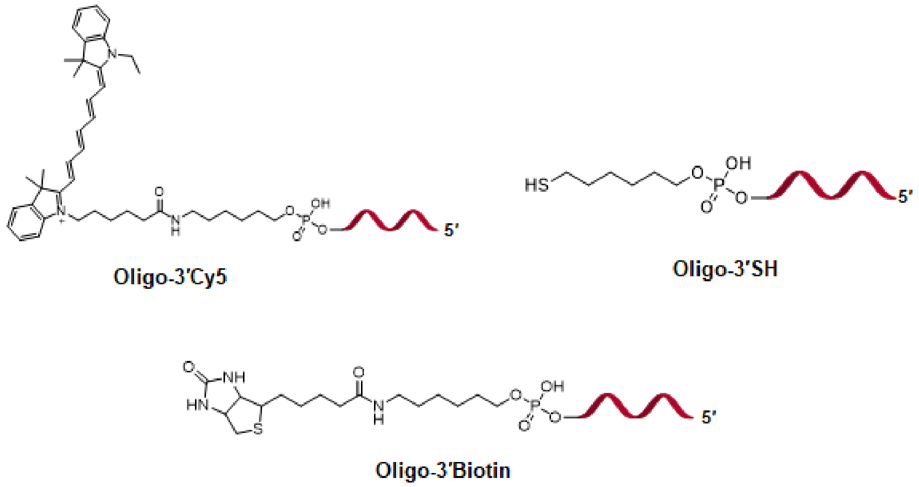


**Figure S5.** Labelled DNAs and PNAs used. **(a)** HPLC and MS characterizations of two PNAs PNA ($NH_2$-Lys-ACCCTAA-$CONH_2$) and PNA ($NH_2$-Lys-CCCTAACCCTAA-$CONH_2$). HPLC Gradient condition: $CH_3CN$ 10% to 50% in 50 min. **(b)** Chemical structures of Cy5, thiol, and biotin groups used for labelling DNAs.

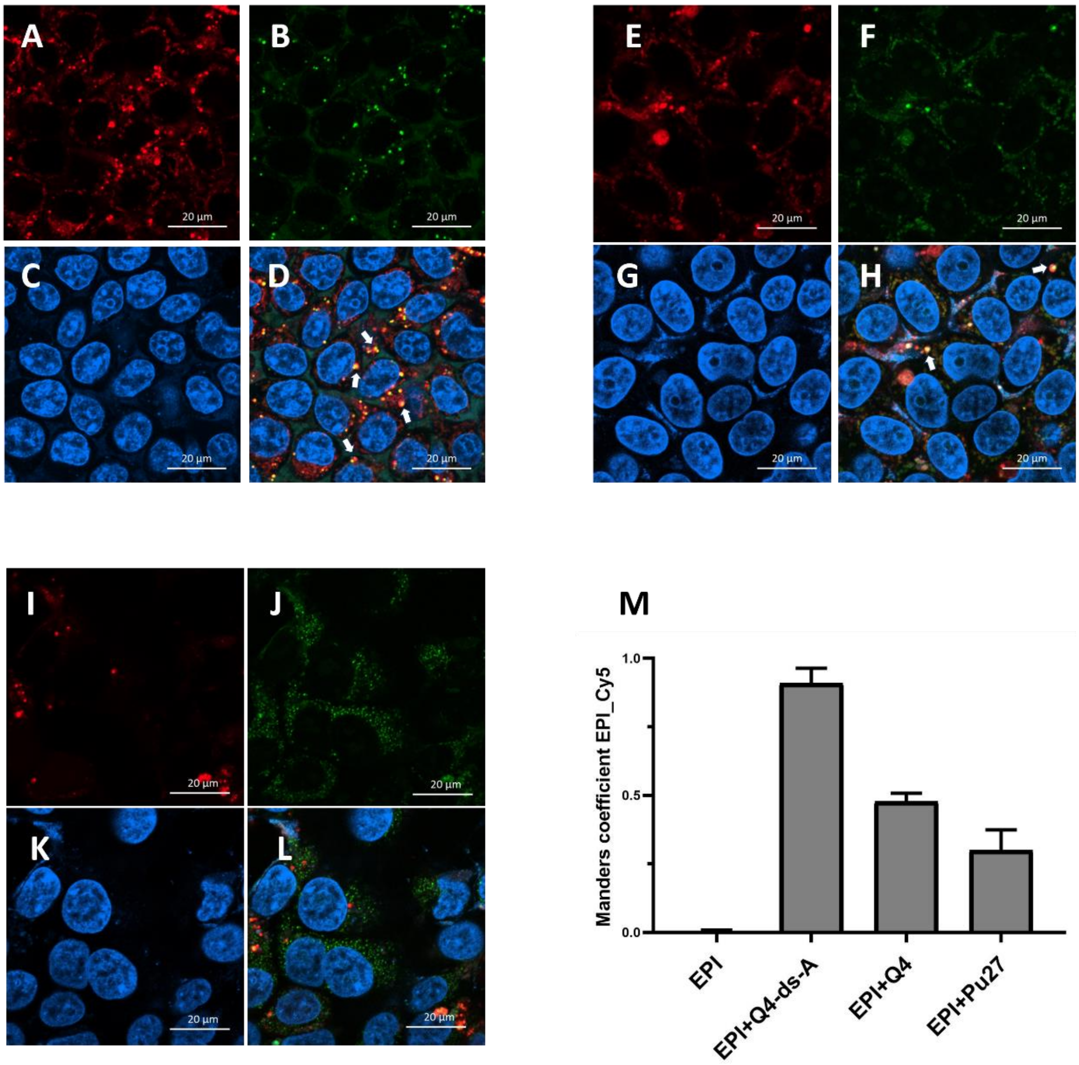


**Figure S6.** Intracellular colocalization assay of EPI with DNA. The red, green and blue fluorescence signals correspond to Cy5-labelled Q4, EPI, and Hoechst, respectively. **(A-D)** Confocal imaging of Cy5-Q4 (1 μM), EPI (10 μM) and Hoechst in HEK293T cells. **(E-H)** Confocal imaging assay results after treatment with Cy5-Q4-ds-A (1 μM), EPI (10 μM) and Hoechst. **(I-L)** Confocal imaging assay results of HEK293T cells treated with Cy5-Pu27 (1 μM), EPI (10 μM) and Hoechst in HEK293T cells. For each of the data sets, (top left, panels **A,E,I**) Cy5 signal, $\lambda_{ex}$ = 640 nm. (top right, panels **B,F,J**) EPI signal, $\lambda_{ex}$ = 488 nm. (bottom left, panels **C,G,K**) Hoechst signal, $\lambda_{ex}$ = 405 nm. (bottom right, panels **D,H,L**) merged image (arrows here show some representative overlapped orange dots). See operational details in the experimental section. **(M)** The Manders coefficient analysis revealed the colocalization extent order: EPI-Cy5-Q4-ds-A > EPI-Cy5-Q4 > EPI-Cy5-Pu27. DNA Cy5-Pu27 was applied as a negative control.

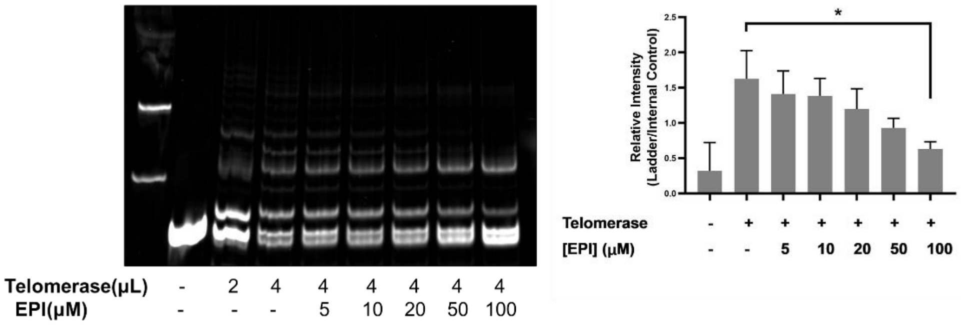


**Figure S7.** TRAP assay of EPI inhibition of telomerase activity. A). TRAP assay for telomerase activity. The internal control (56 bp) is indicated in the left lane. The length of the shortest band in the ladder is 61 bp. The telomerase products formed a 6 bp-ladder in each lane. B) Quantitative analysis of the ratio of intensity levels of the telomerase products and internal control after background subtraction. ImageJ software was used to determine the intensity. The experiments were conducted three times.

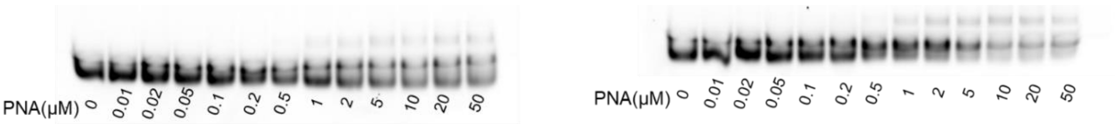


**Figure S8.** Nondenaturing PAGE of a 7-mer PNA binding to DNA. Titration of PNA ($NH_2$-Lys-ACCCTAA-$CONH_2$) into Q4 with (*left*) and without (*right*) the addition of 2 μM EPI. The samples were prepared by snap cooling of the hairpins (heated at 95 °C for 5 min, then immediately put on ice and equilibrated at 4 °C for 10 min), followed by annealing with the PNA oligomer by heating at 65 °C for 5 min followed by slow cooling to room temperature for about 2 hours and incubation at 4 °C about 20 hours. Samples were incubated with 25 mM Tris-HCl, 100mM KCl, 1mM EDTA, pH=7.5.

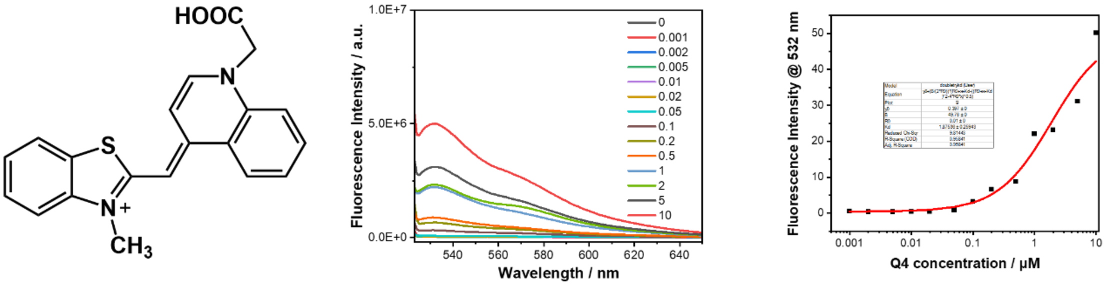


**Figure S9.** $K_D$ determination via fluorescence titrating assay using varied DNA **Q4** concentrations 0-10 μM and 10 nM TO, in 110 μl of a buffer consisting of 100 mM KCl, 0.5 mM EDTA, and 20 mM tris (pH 7.5). All of the emission spectra were recorded at room temperature, at an excitation wavelength of 377 nm. Fitting $R^2$ = 0.95 and $K_D$ is **1870 ± 260 nM**. The chemical structure of TO (Thiazole orange) is shown in the left panel.

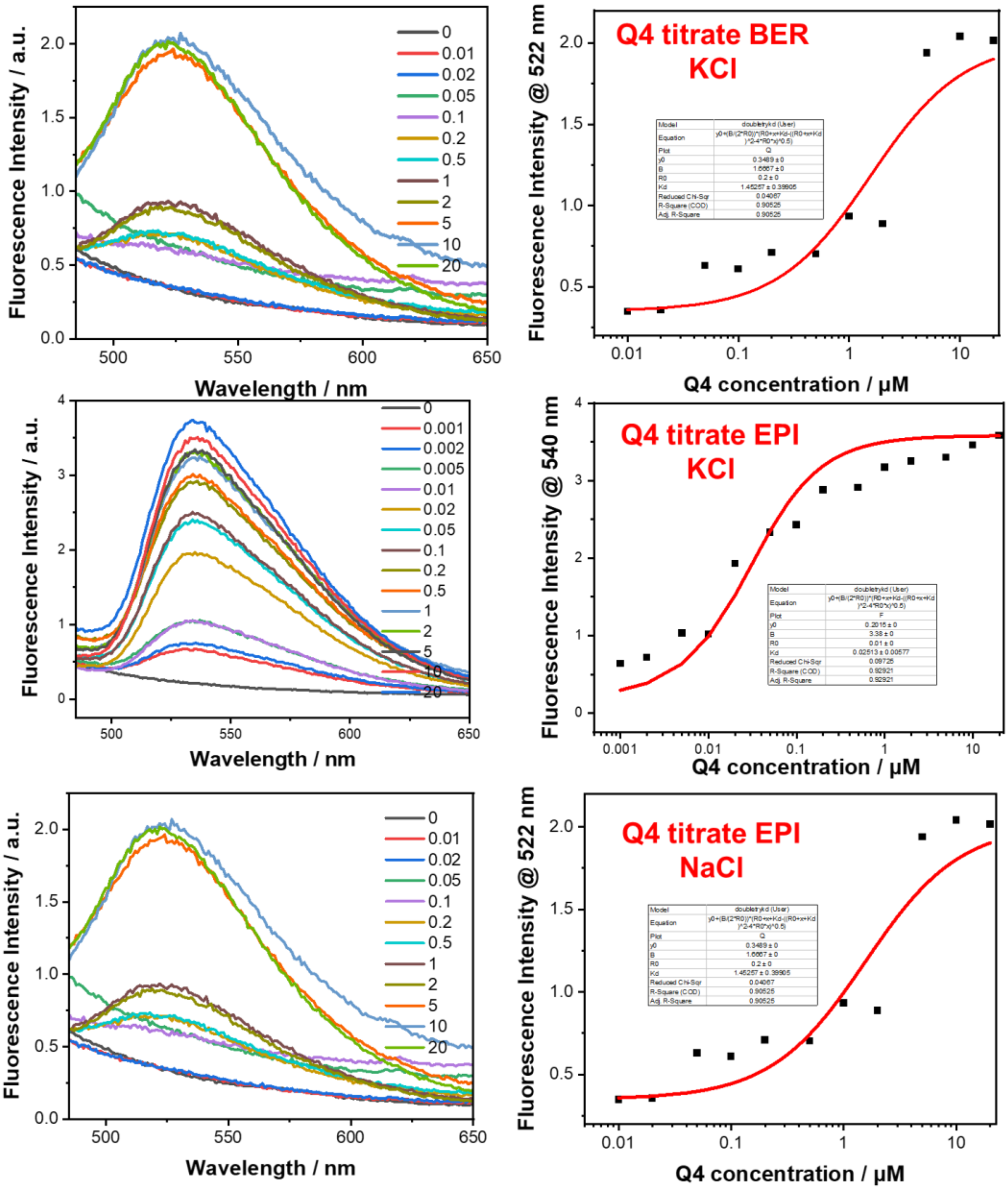


**Figure S10.** $K_D$ determination via fluorescence titrating assay using varied DNA **Q4** concentrations 0-10 µM and 10 nM BER or EPI, in 110 µl of a buffer consisting of 100 mM KCl or 100 mM NaCl, 0.5 mM EDTA, and 20 mM tris (pH 7.5). All of the emission spectra were recorded at room temperature, at an excitation wavelength of 377 nm. (Top) For Q4 titrated into BER in KCl buffer: fitting $R^2$ = 0.90 and $K_D$ is **1450 ± 390 nM**. (Middle) For Q4 titrated into EPI in KCl buffer: fitting $R^2$ = 0.93 and $K_D$ is **26 ± 4 nM**. (Bottom) For Q4 titrated into EPI in NaCl buffer: fitting $R^2$ = 0.90 and $K_D$ is **1379 ± 198 nM**.

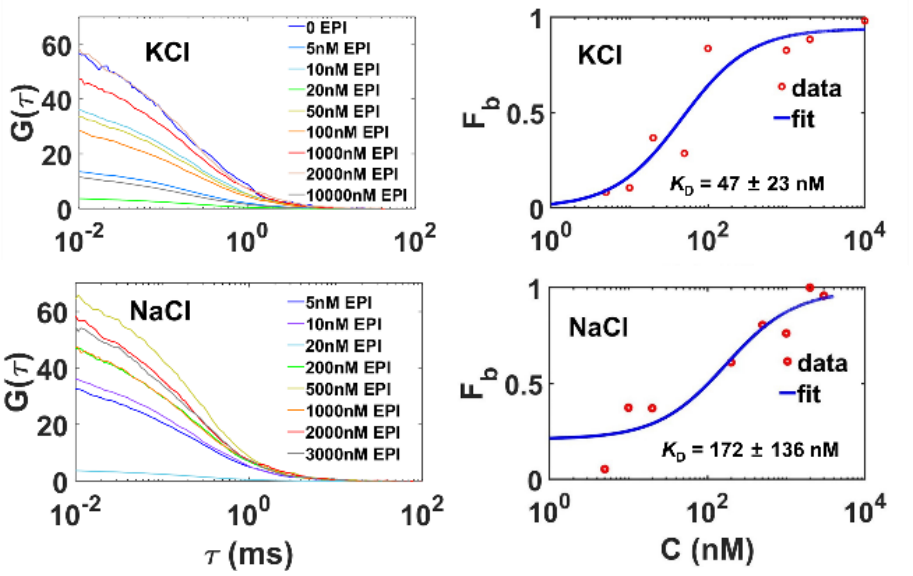


**Figure S11.** Competitive binding of EPI and PNA to Cy5-labeled Q4 characterization by FCS. The Cy5 fluorescence autocorrelation curves were obtained for the samples composed of 1 nM Cy5-Q4 (with Cy5 covalently attached on Q4), 100 nM 12-mer PNA ($NH_2$-lys-CCCTAACCCTAA-$CONH_2$), and different concentrations of EPI (0-10 μM) in 0.1 M KCl **(Top left)** or 0.1 M NaCl **(Bottom left)**. $K_D$ values of EPI competitive binding to Cy5-Q4 were obtained by nonlinear least squares fitting of the binding data in 0.1 M KCl **(Top right)** or 0.1 M NaCl **(Bottom right)**.

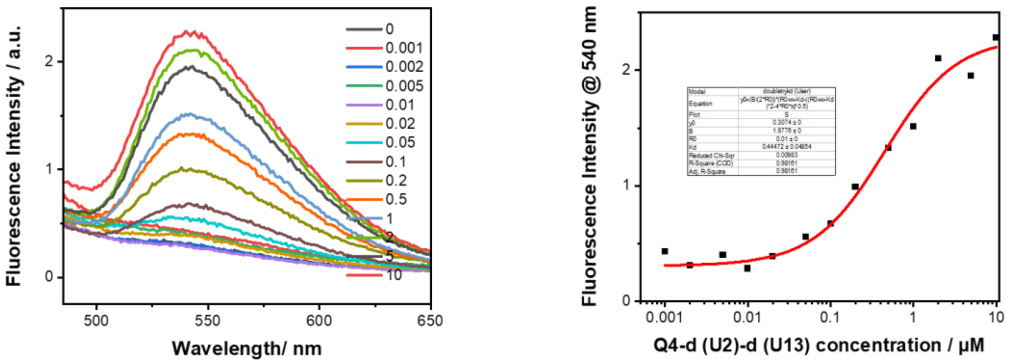


**Figure S12.** $K_D$ determination via fluorescence titrating assay using varied DNA **Q4-d(U2)-d(U13)** concentrations 0-10 μM and 10 nM EPI, in 110 μl of a buffer consisting of 100 mM KCl, 0.5 mM EDTA, and 20 mM tris (pH 7.5). All of the emission spectra were recorded at room temperature, at an excitation wavelength of 377 nm. Fitting $R^2$ = 0.98 and $K_D$ is **444 ± 48 nM**.

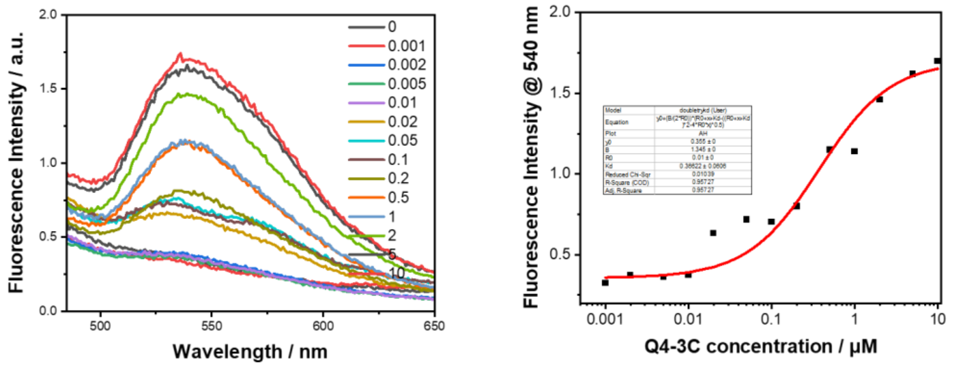


**Figure S13.** $K_D$ determination via fluorescence titrating assay using varied DNA **Q4-3C** concentrations 0-10 μM and 10 nM EPI, in 110 μl of a buffer consisting of 100 mM KCl, 0.5 mM EDTA, and 20 mM tris (pH 7.5). All of the emission spectra were recorded at room temperature, at an excitation wavelength of 377 nm. Fitting $R^2$ = 0.95 and $K_D$ is **366 ± 61 nM**.

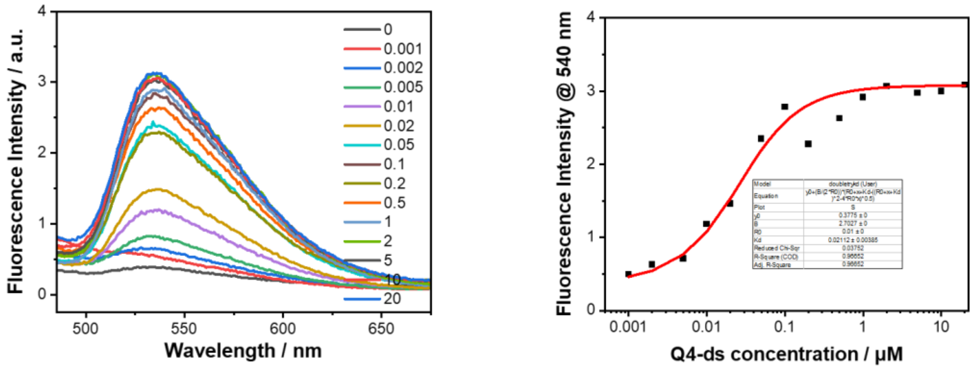


**Figure S14.** $K_D$ determination via fluorescence titrating assay using varied DNA **Q4-ds** concentrations 0-20 μM and 10 nM EPI, in 110 μl of a buffer consisting of 100 mM KCl, 0.5 mM EDTA, and 20 mM tris (pH 7.5). All of the emission spectra were recorded at room temperature, at an excitation wavelength of 377 nm. Fitting $R^2$ = 0.96 and $K_D$ is **21 ± 4 nM**.

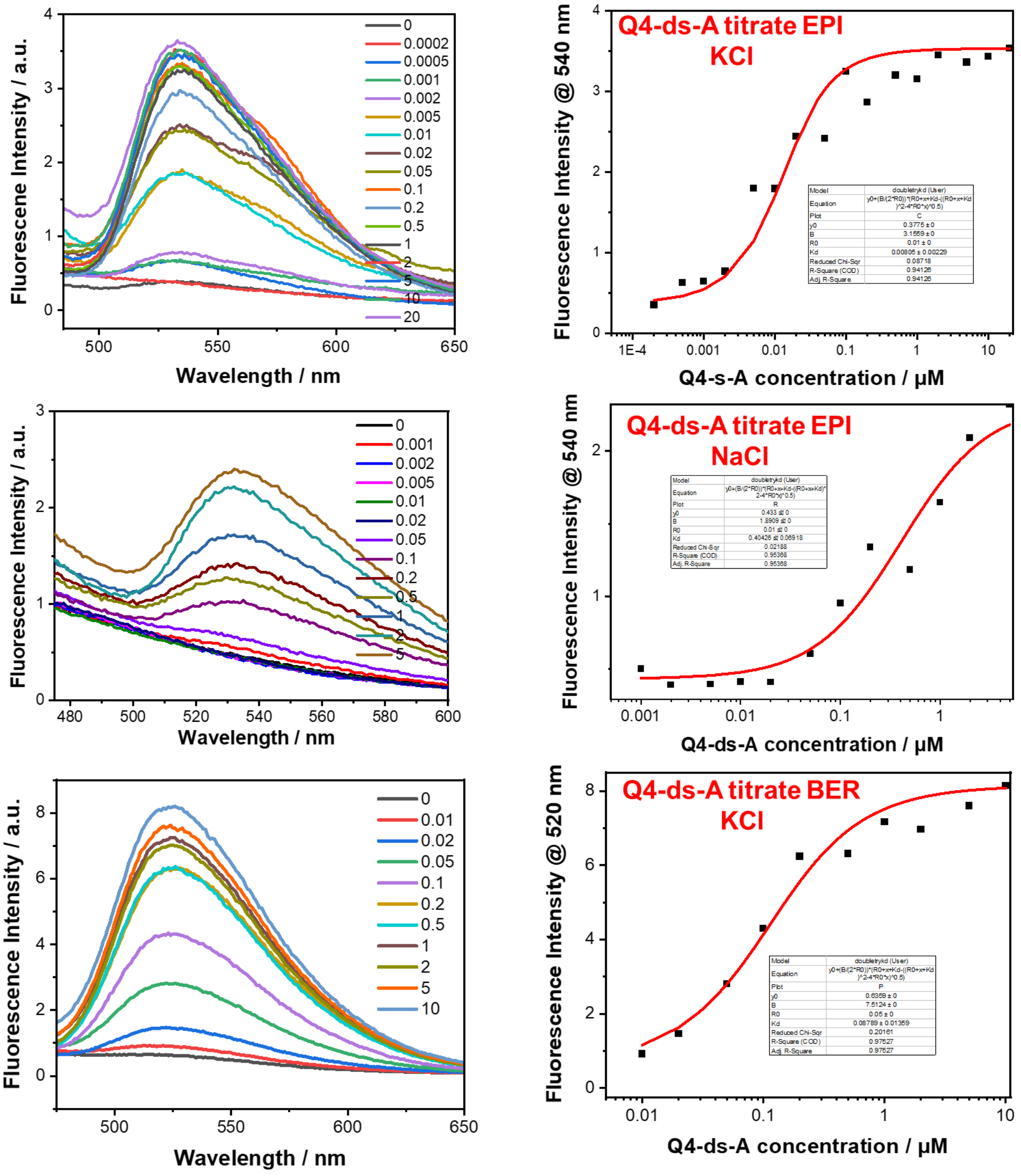


**Figure S15.** $K_D$ determination via fluorescence titrating assay using varied DNA **Q4-ds-A** concentrations 0-20 µM and 10 nM EPI or BER, in 110 µl of a buffer consisting of 100 mM KCl/ NaCl, 0.5 mM EDTA, and 20 mM tris (pH 7.5). All of the emission spectra were recorded at room temperature, at an excitation wavelength of 377 nm. For Q4-ds-A titrated into EPI in KCl: fitting $R^2 = 0.94$ and $K_D$ is **8 ± 2 nM**. For Q4-ds-A titrated into EPI in NaCl: fitting $R^2 = 0.95$ and $K_D$ is **400±69 nM**. For Q4-ds-A titrated into BER in KCl: fitting $R^2 = 0.97$ and $K_D$ is **88±13 nM**.

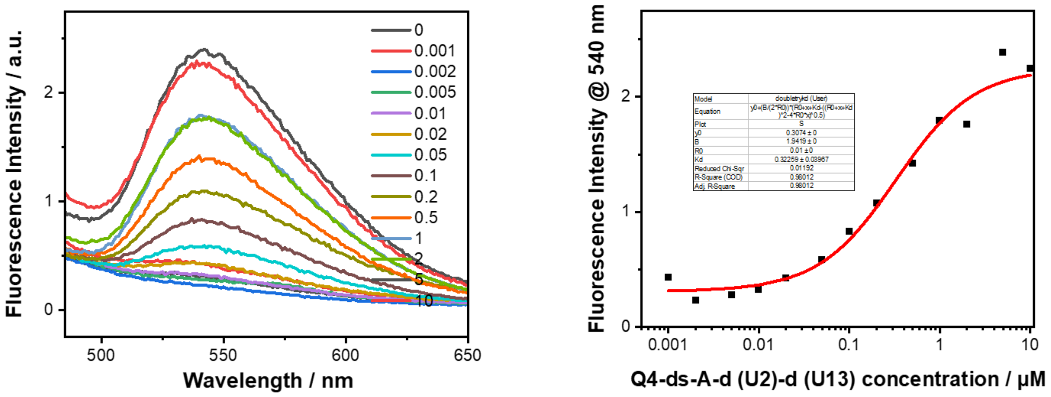


**Figure S16.** $K_D$ determination via fluorescence titrating assay using varied DNA **Q4-ds-A-d(U2)-d(U13)** concentrations 0-10 μM and 10 nM EPI, in 110 μl of a buffer consisting of 100 mM KCl, 0.5 mM EDTA, and 20 mM tris (pH 7.5). All of the emission spectra were recorded at room temperature, at an excitation wavelength of 377 nm. Fitting $R^2$ = 0.98 and $K_D$ is **322 ± 39 nM**.

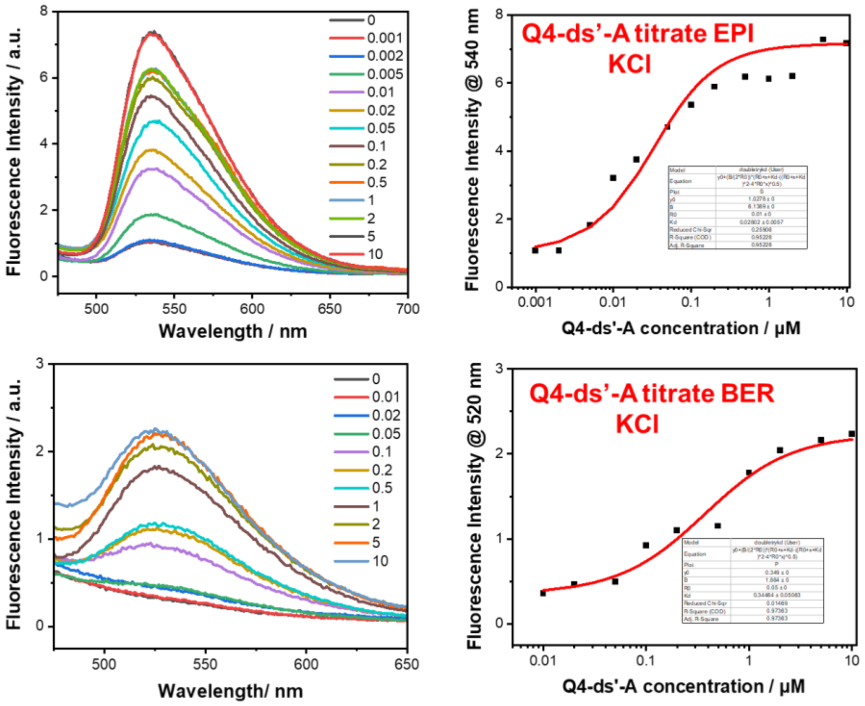


**Figure S17**. (Top) $K_D$ determination via fluorescence titrating assay using varied DNA **Q4-ds′-A** concentrations 0-10 μM and 10 nM EPI, in 110 μl of a buffer consisting of 100 mM KCl, 0.5 mM EDTA, and 20 mM tris (pH 7.5). All of the emission spectra were recorded at room temperature, at an excitation wavelength of 377 nm. Fitting $R^2$ = 0.95 and $K_D$ is **28 ± 6 nM**. (Bottom) $K_D$ determination via fluorescence titrating assay using varied DNA **Q4-ds′-A** concentrations 0-10 μM and 10 nM BER, in 110 μl of a buffer consisting of 100 mM KCl, 0.5 mM EDTA, and 20 mM tris (pH 7.5). All of the emission spectra were recorded at room temperature, at an excitation wavelength of 377 nm. Fitting $R^2$ = 0.97 and $K_D$ is **344 ± 51 nM**.

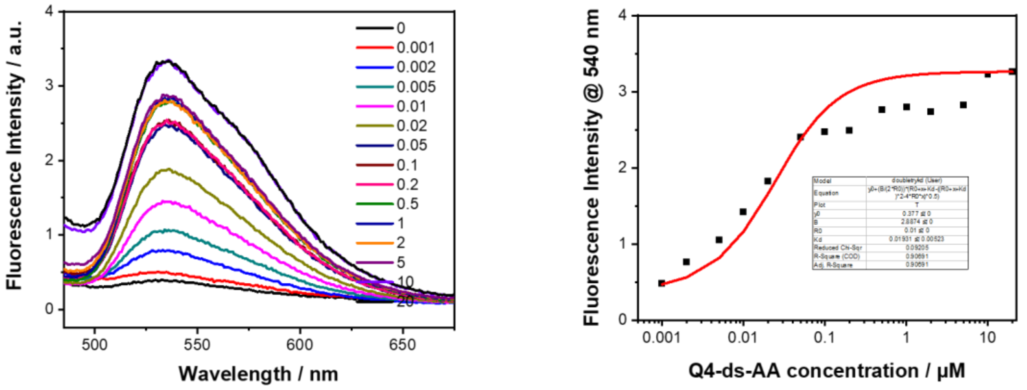


**Figure S18.** $K_D$ determination via fluorescence titrating assay using varied DNA **Q4-ds-AA** concentrations 0-20 μM and 10 nM EPI, in 110 μl of a buffer consisting of 100 mM KCl, 0.5 mM EDTA, and 20 mM tris (pH 7.5). All of the emission spectra were recorded at room temperature, at an excitation wavelength of 377 nm. Fitting $R^2 = 0.9$ and $K_D$ is **19 ± 5 nM**.

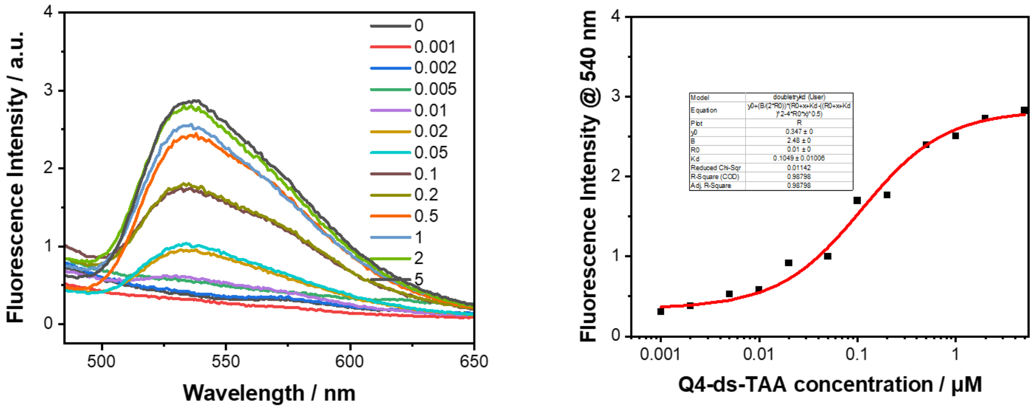


**Figure S19.** $K_D$ determination via fluorescence titrating assay using varied DNA **Q4-ds-TAA** concentrations 0-5 μM and 10 nM EPI, in 110 μl of a buffer consisting of 100 mM KCl, 0.5 mM EDTA, and 20 mM tris (pH 7.5). All of the emission spectra were recorded at room temperature, at an excitation wavelength of 377 nm. Fitting $R^2 = 0.98$ and $K_D$ is **105 ± 10 nM**.

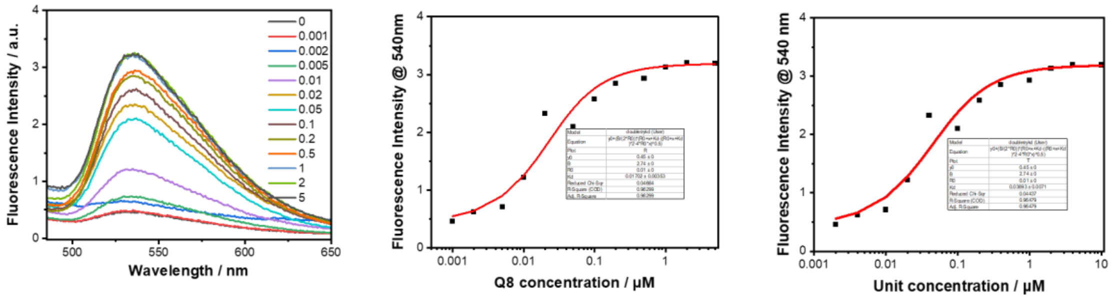


**Figure S20.** $K_D$ determination via fluorescence titrating assay using varied DNA **Q8** concentrations 0-5 µM and 10 nM EPI, in 110 µl of a buffer consisting of 100 mM KCl, 0.5 mM EDTA, and 20 mM tris (pH 7.5). All of the emission spectra were recorded at room temperature, at an excitation wavelength of 377 nm. Fitting $R^2$ = 0.96 and $K_D$ is **17 ± 4 nM** (DNA strand concentration in X-axis). Fitting $R^2$ = 0.96 and $K_D$ is **39 ± 7 nM** (G-quadruplex unit concentration in X-axis).

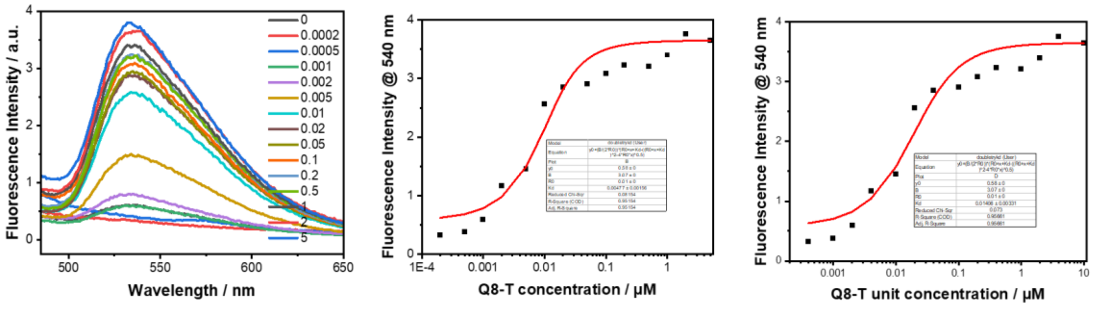


**Figure S21.** $K_D$ determination via fluorescence titrating assay using varied DNA **Q8-T** concentrations 0-5 µM and 10 nM EPI, in 110 µl of a buffer consisting of 100 mM KCl, 0.5 mM EDTA, and 20 mM tris (pH 7.5). All of the emission spectra were recorded at room temperature, at an excitation wavelength of 377 nm. Fitting $R^2$ = 0.95 and $K_D$ is **5 ± 2 nM** (DNA strand concentration in X-axis). Fitting $R^2$ = 0.95 and $K_D$ is **14 ± 3 nM** (G-quadruplex unit concentration in X-axis).

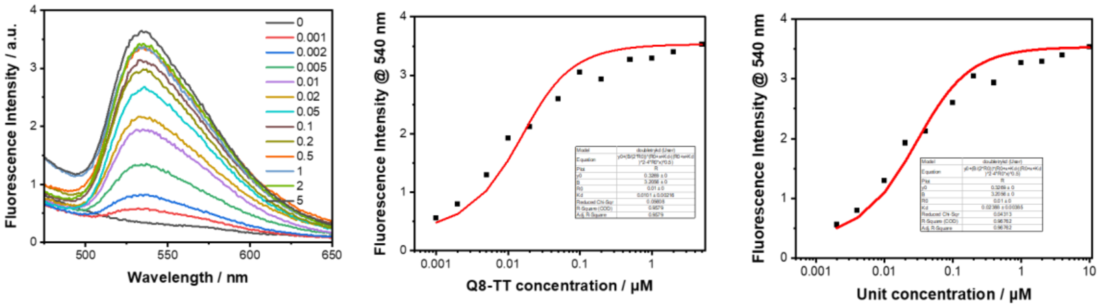


**Figure S22.** $K_D$ determination via fluorescence titrating assay using varied DNA **Q8-TT** concentrations 0-5 µM and 10 nM EPI, in 110 µl of a buffer consisting of 100 mM KCl, 0.5 mM EDTA, and 20 mM tris (pH 7.5). All of the emission spectra were recorded at room temperature, at an excitation wavelength of 377 nm. Fitting $R^2$ = 0.95 and $K_D$ is **10 ± 2 nM** (DNA concentration as x-axis). Fitting $R^2$ = 0.96 and $K_D$ is **24 ± 4 nM** (unit concentration as x-axis).

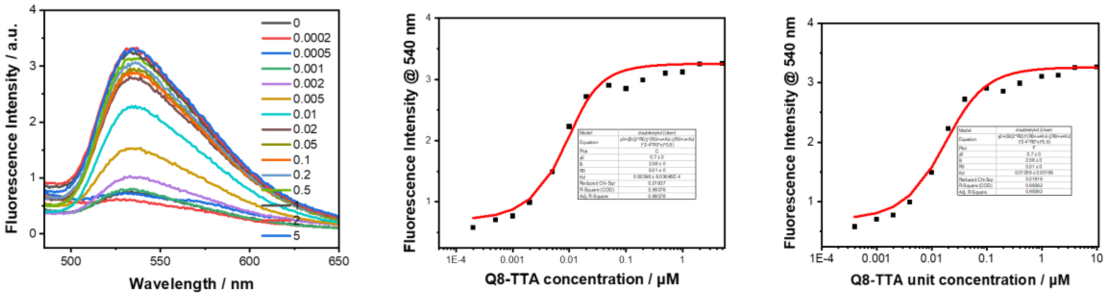


**Figure S23.** $K_D$ determination via fluorescence titrating assay using varied DNA **Q8-TTA** concentrations 0-5 μM and 10 nM EPI, in 110 μl of a buffer consisting of 100 mM KCl, 0.5 mM EDTA, and 20 mM tris (pH 7.5). All of the emission spectra were recorded at room temperature, at an excitation wavelength of 377 nm. Fitting $R^2$ = 0.98 and $K_D$ is **4 ± 1 nM** (DNA concentration as x-axis). Fitting $R^2$ = 0.98 and $K_D$ is **13 ± 2 nM** (G-quadruplex unit concentration in X-axis).

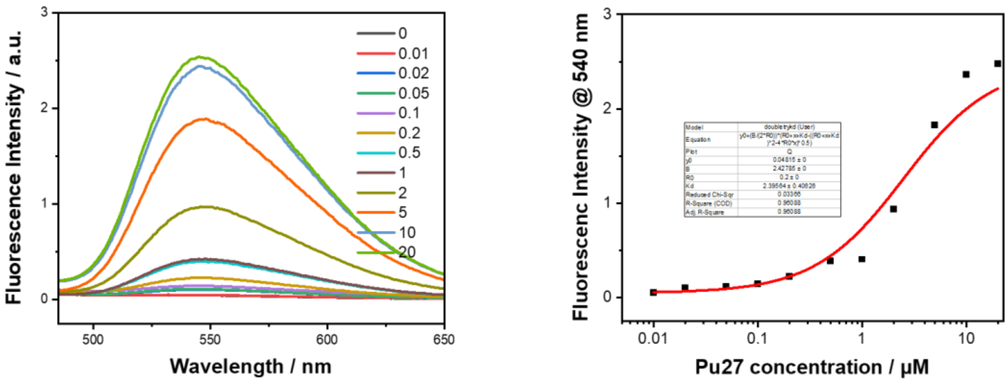


**Figure S24.** $K_D$ determination via fluorescence titrating assay using varied DNA **Pu27** (5'-TGGGGAGGGTGGGGAGGGTGGGGAAGG-3') concentrations 0-20 μM and **0.2 μM** EPI, in 110 μl of a buffer consisting of 100 mM KCl, 0.5 mM EDTA, and 20 mM tris (pH 7.5). All of the emission spectra were recorded at room temperature, at an excitation wavelength of 377 nm. Fitting $R^2$ = 0.96 and $K_D$ is **2390 ± 400 nM**.

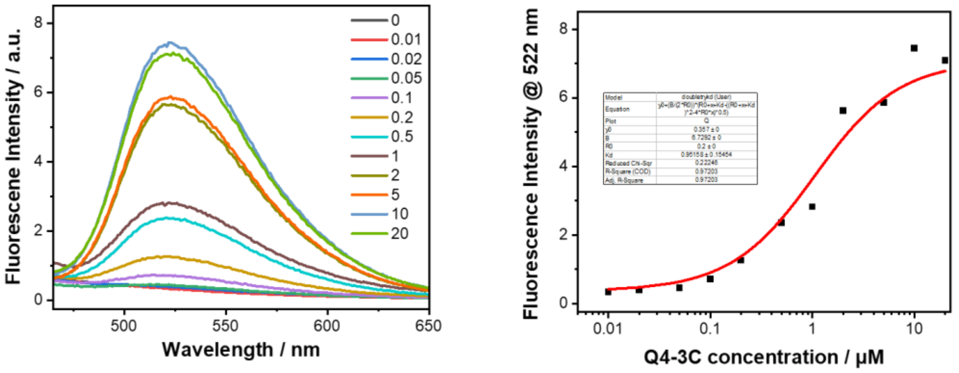


**Figure S25.** $K_D$ determination via fluorescence titrating assay using varied DNA **Q4-3C** concentrations 0-20 μM and **0.2 μM BER**, in 110 μl of a buffer consisting of 100 mM KCl, 0.5 mM EDTA, and 20 mM tris (pH 7.5). All of the emission spectra were recorded at room temperature, at an excitation wavelength of 377 nm. Fitting $R^2$ = 0.97 and $K_D$ is **950 ± 150 nM**.

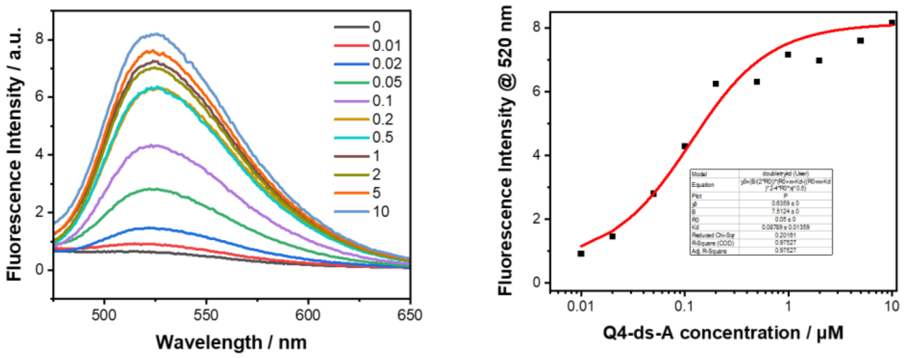


**Figure S26.** $K_D$ determination via fluorescence titrating assay using varied DNA **Q4-ds-A** concentrations 0-10 μM and **50 nM BER**, in 110 μl of a buffer consisting of 100 mM KCl, 0.5 mM EDTA, and 20 mM tris (pH 7.5). All of the emission spectra were recorded at room temperature, at an excitation wavelength of 377 nm. Fitting $R^2 = 0.97$ and $K_D$ is **88 ± 13 nM**.

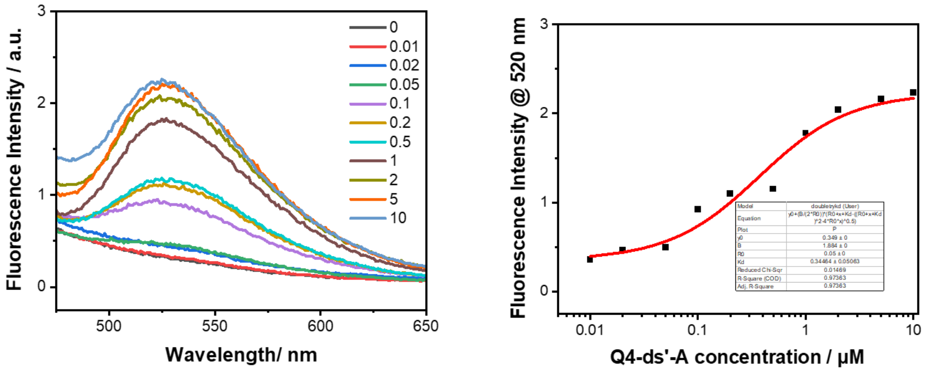


**Figure S27.** $K_D$ determination via fluorescence titrating assay using varied DNA **Q4-ds′-A** concentrations 0-10 μM and **50 nM BER**, in 110 μl of a buffer consisting of 100 mM KCl, 0.5 mM EDTA, and 20 mM tris (pH 7.5). All of the emission spectra were recorded at room temperature, at an excitation wavelength of 377 nm. Fitting $R^2 = 0.97$ and $K_D$ is **344 ± 51 nM**.

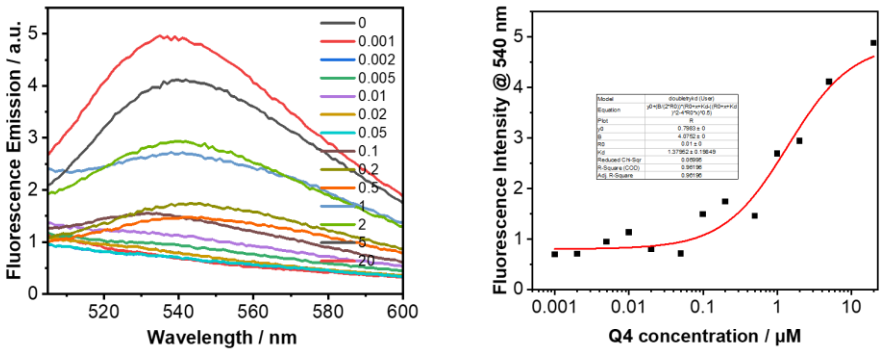


**Figure S28.** $K_D$ determination via fluorescence titrating assays using varied DNA **Q4** concentration 0-20 μM and 10 nM EPI, in 110 μl of a buffer consisting of 100 mM **NaCl**, 0.5 mM EDTA, and 20 mM tris (pH 7.5). All of the emission spectra were recorded at room temperature, at an excitation wavelength of 377 nm. Fitting $R^2 = 0.95$ and $K_D$ is**1379 ± 198 nM**.

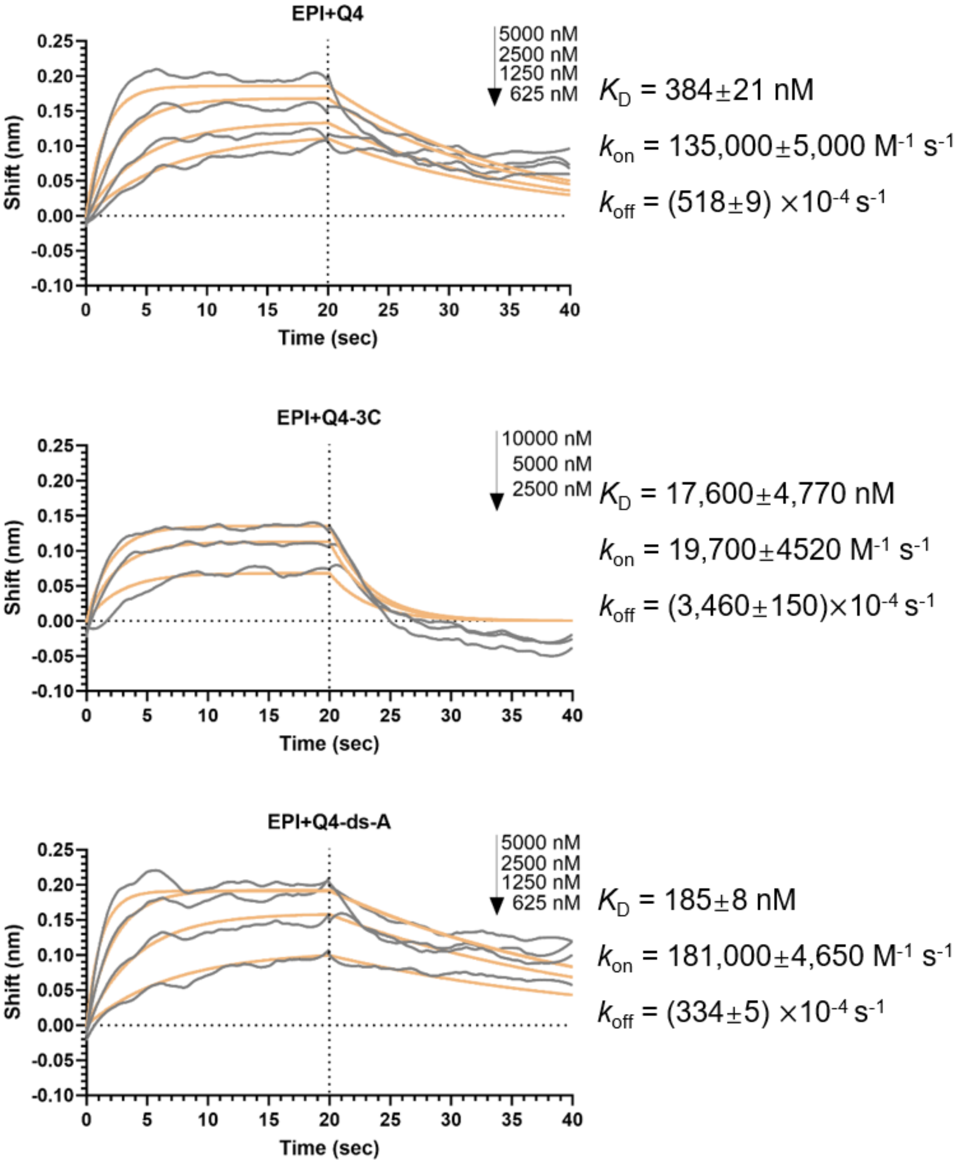


**Figure S29.** Binding kinetics measurement by bio-layer interferometry (BLI). The binding buffer consists of 100 mM KCl, 0.5 mM EDTA, and 20 mM tris (pH 7.5). The measurements were carried out at 30°C.

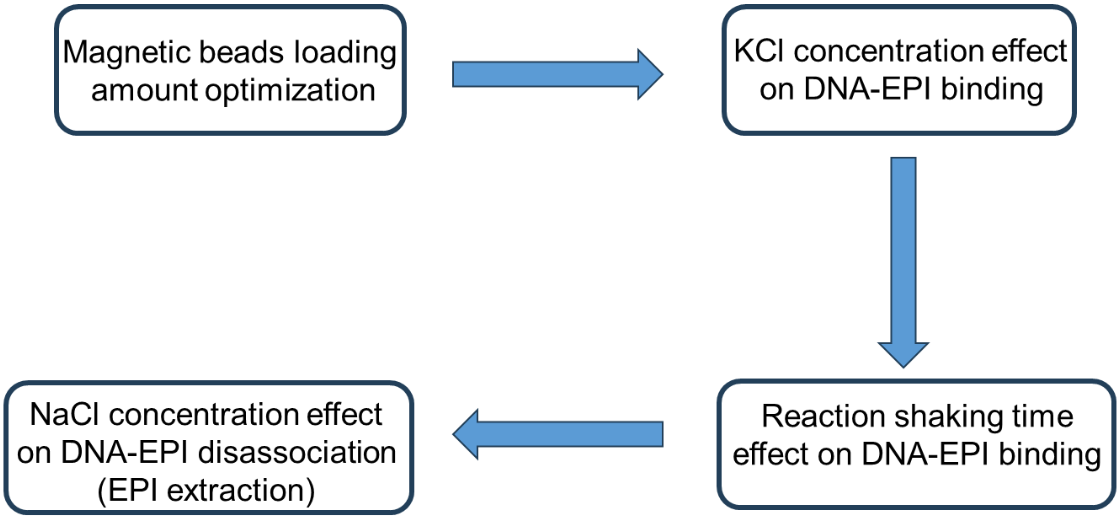


**Scheme S1.** Optimization experiments conducted for the recovery of EPI from a pure EPI solution.

**Probe the magnetic beads loading amount**

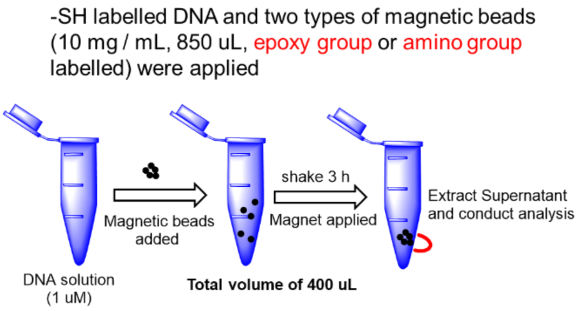


**Measured by Nanodrop**

| Entry | Content | A260 (DNA solution) | A260 (Supernatant) |
|---|---|---|---|
| 1 | Q4-SH + epoxy-MB (5 uL) | 0.14 | 0.04 |
| 2 | Q4-SH + epoxy-MB (10 uL) | 0.14 | 0 |
| 3 | Q4-SH + amino-MB (5 uL) | 0.14 | 0.03 |
| 4 | Q4-SH + amino-MB (10 uL) | 0.14 | 0 |

**10 uL MB (from stocking solution) is enough for coupling with –SH DNA (1 uM, 400 uL)**

**All used KCl and NaCl buffer contained 0.5 mM EDTA, 20 mM Tris, pH 7.5**

**Figure S30.** Optimization of magnetic beads loading.

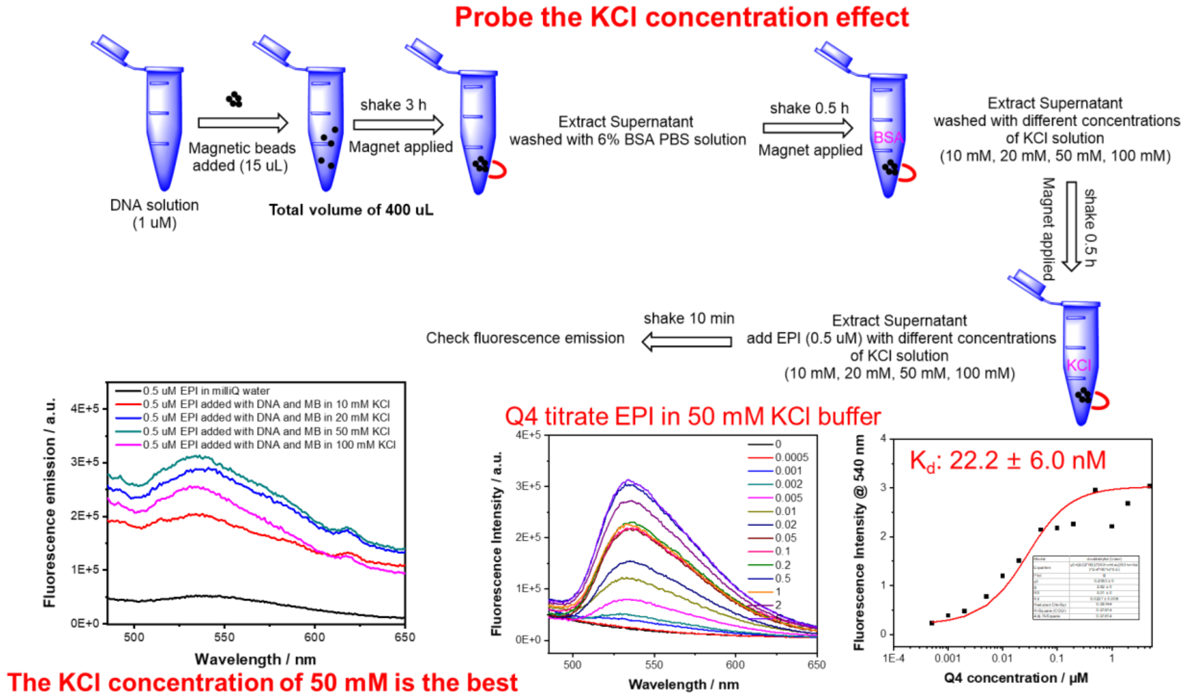


**Figure S31.** Effect of KCl concentration on EPI binding to Q4.

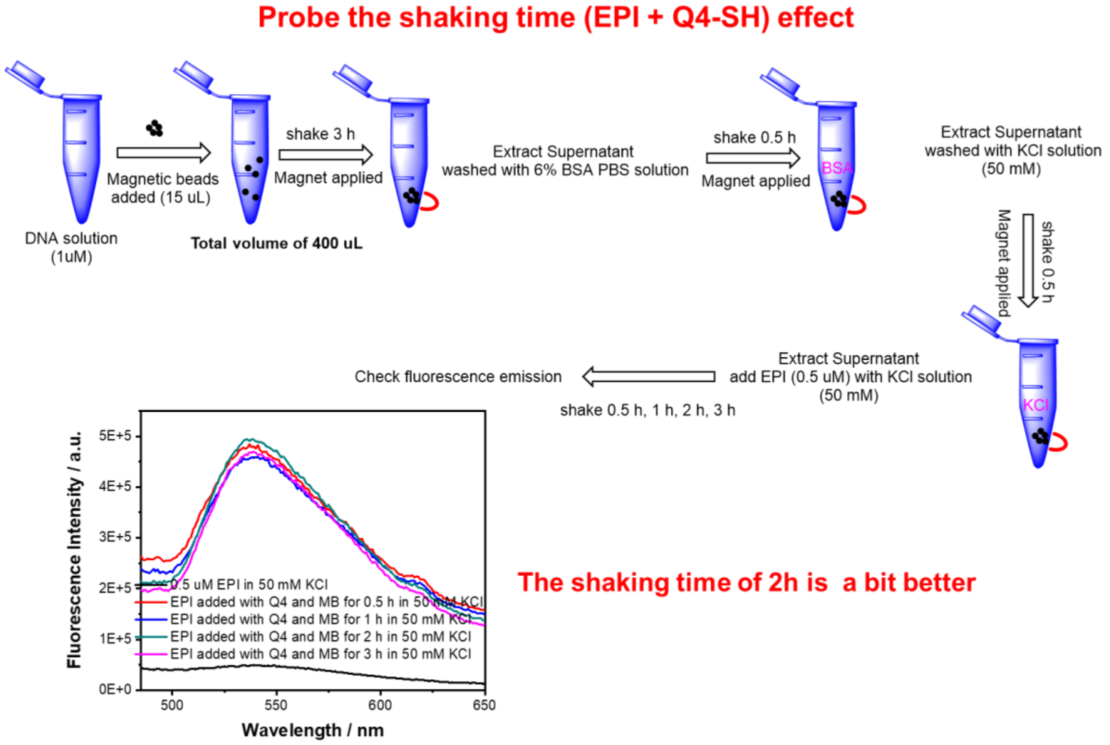


**Figure S32.** Effect of incubation time on EPI binding to Q4.

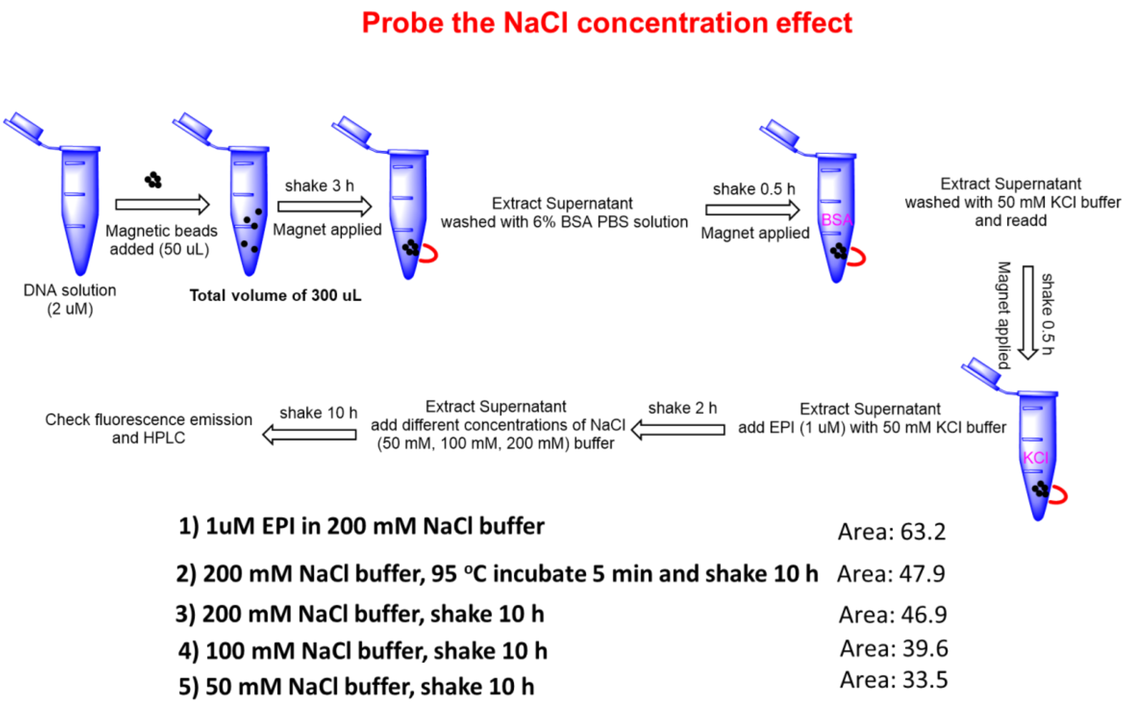


**Figure S33.** Effect of NaCl concentration on EPI release from Q4. See the HPLC data in Figure S32.

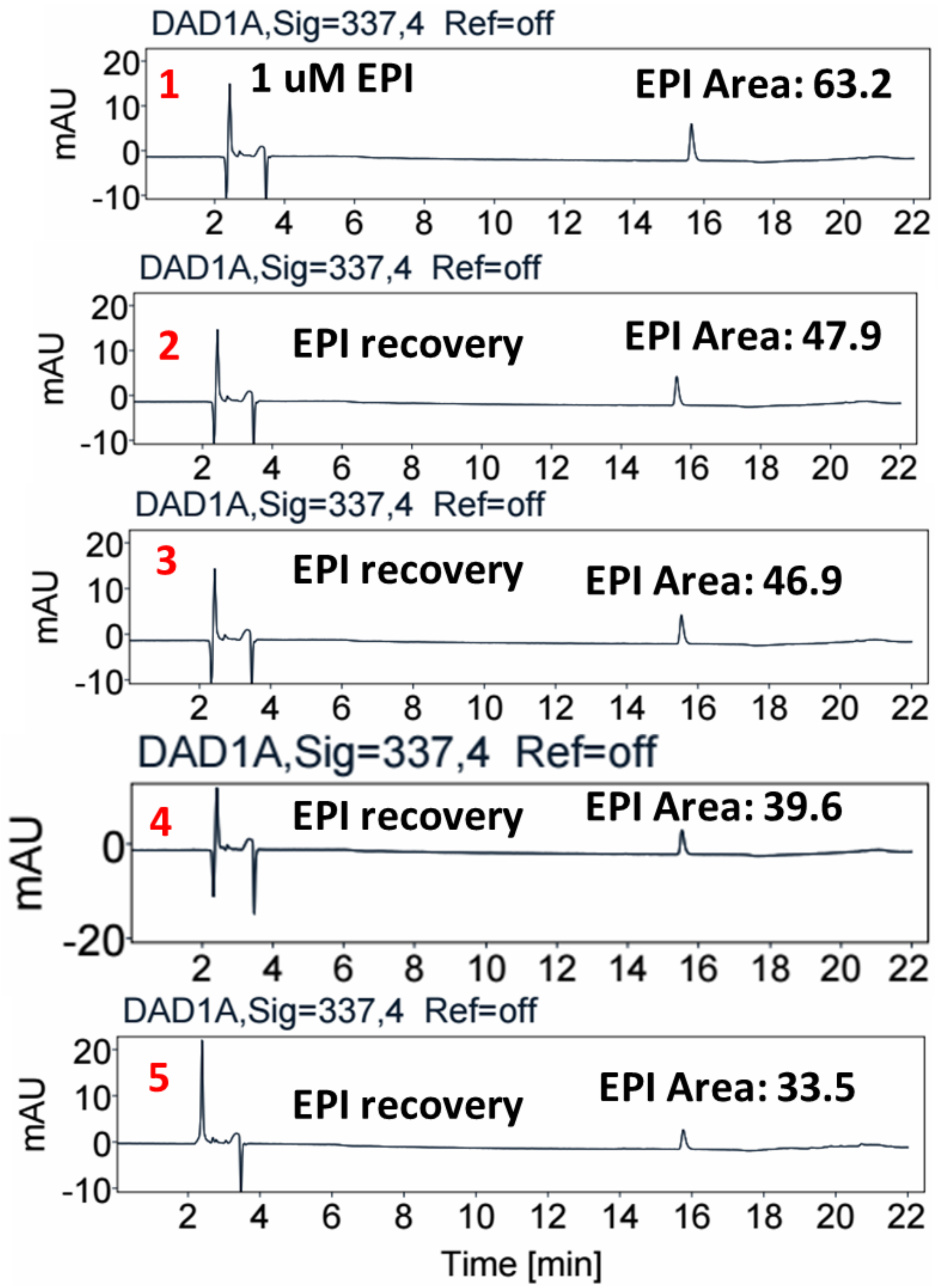


**Figure S34.** HPLC traces of entries 1-5 stated in **Figure S31**.

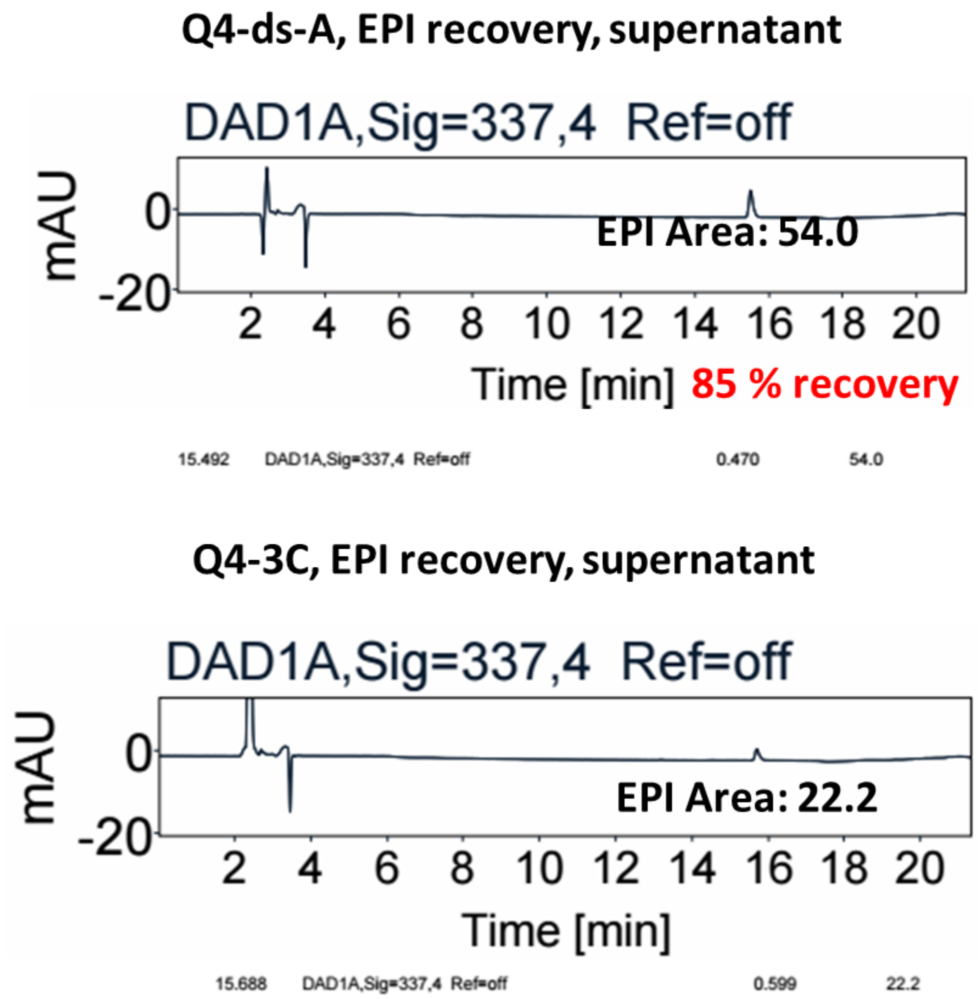


**Figure S35.** HPLC characterization of EPI recovery from a pure EPI solution by using 50 mM KCl for EPI binding followed by 200 mM NaCl for EPI release. Q4-ds-A shows a higher EPI recovery rate (85%)compared with Q4-3C (34%).

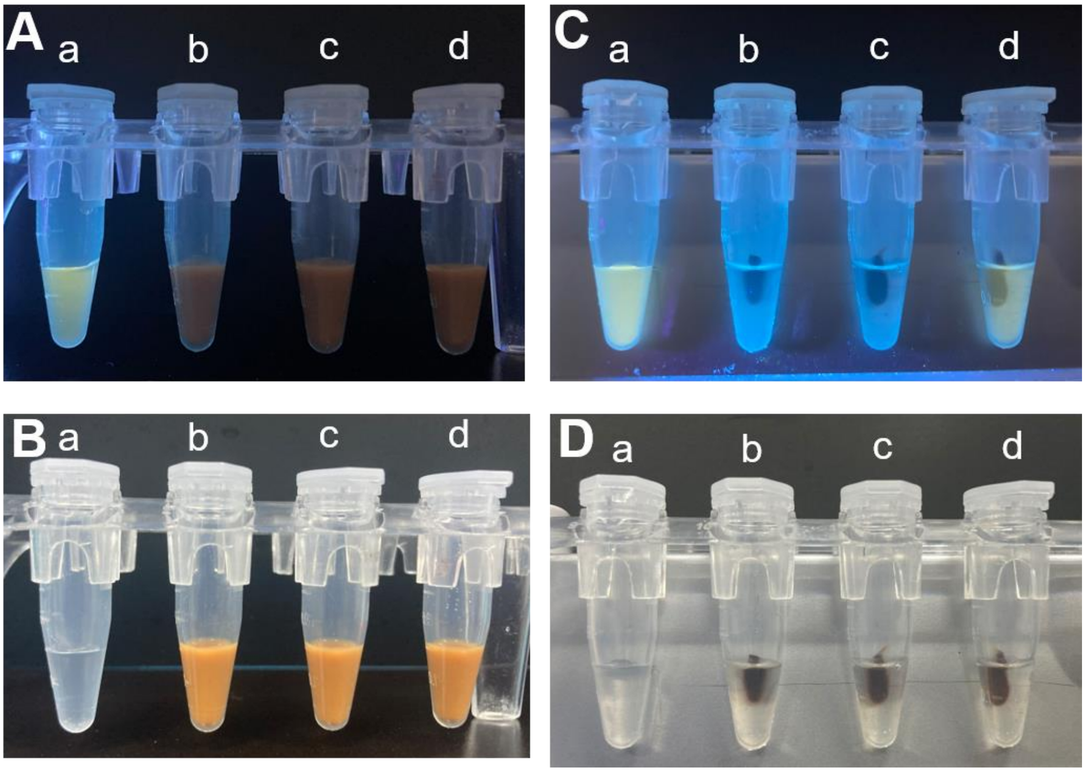


**Figure S36.** Visualization of fluorescence light-up effect of EPI by binding to Q4-ds-A. The photos were taken without (A,B) and with (C,D) a magnet applied. The photos were taken under UV (365 nm, A,C) and room light (B,D). In each panel, the test tubes contain (a) thiolated Q4-ds-A and EPI, (b) EPI with MB, (c) EPI and MB labelled with thiolated Q4-ds-A, and (d) EPI, nonthiolated Q4-ds-A and MB. The DNA and EPI are at 1 μM in a solution of 50 mM KCl. The MB is at 10 mg/ml.

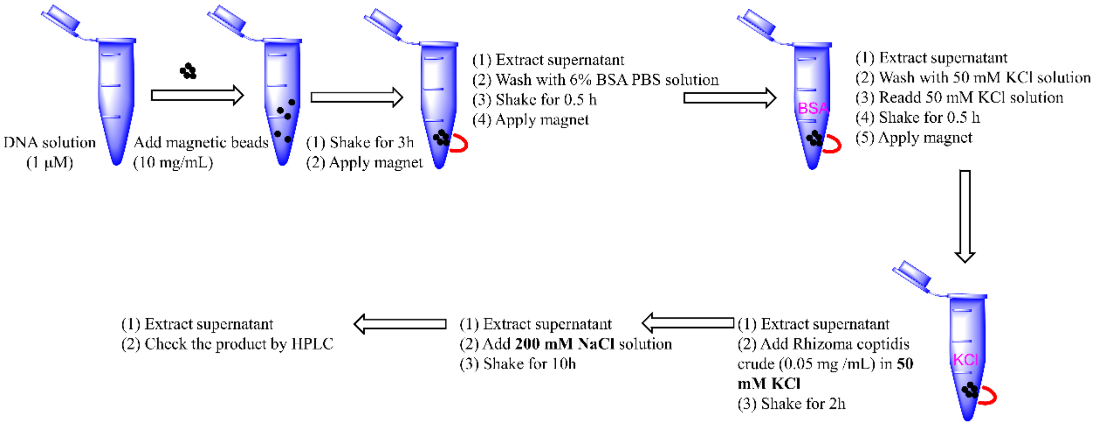


**Figure S37.** Optimized protocol for EPI affinity extraction from the *Rhizoma coptidis* (Huang Lian) extracts.

**Table S1.** Summary of HPLC characterization data for the EPI extraction protocol. For the first three batches, a buffer containing 200 mM NaCl was used for EPI release. For the fourth batch, a solution of diluted HCl (pH 3) was used for EPI release.

| HPLC peak area | | | supernatant of 50 mM KCl buffer treatment | | | Supernatant of 200 mM NaCl or diluted HCl treatment | | | EPI recovery percentage (%) |
|---|---|---|---|---|---|---|---|---|---|
| | Compound | Crude | Q4-ds-A | Q4 | Q4-3C | Q4-ds-A | Q4 | Q4-3C | 57.8 |
| **First batch (EPI elution by NaCl)** | **COL** | 111 | 93 | 97 | 104 | - | - | - | |
| | **EPI** | 729 | 293 | 374 | 637 | 422 | 301 | 11 | |
| | **COP** | 360 | 280 | 299 | 333 | - | 20 | 3 | |
| | **PAL** | 511 | 409 | 437 | 477 | 26 | 26 | 4 | |
| | **BER** | 2111 | 1666 | 1987 | 1948 | 171 | 104 | 19 | |
| **Second batch (EPI elution by NaCl)** | **COL** | 109 | 120 | 128 | 100 | - | - | - | 58.9 |
| | **EPI** | 716 | 332 | 418 | 624 | 422 | 304 | 80 | |
| | **COP** | 359 | 308 | 331 | 332 | - | 22 | 24 | |
| | **PAL** | 483 | 382 | 415 | 455 | 23 | 24 | 26 | |
| | **BER** | 2139 | 1674 | 1947 | 1951 | 171 | 108 | 178 | |
| **Third batch (EPI elution by NaCl)** | **COL** | 125 | 137 | 146 | 157 | - | - | - | 55.1 |
| | **EPI** | 822 | 375 | 472 | 776 | 486 | 329 | 93 | |
| | **COP** | 418 | 363 | 394 | 446 | - | - | 27 | |
| | **PAL** | 587 | 459 | 489 | 531 | 30 | 24 | 29 | |
| | **BER** | 2436 | 1912 | 2236 | 2248 | 208 | 106 | 158 | |
| **Fourth batch (EPI elution by HCl (pH 3))** | **COL** | 155 | 118 | - | - | - | - | - | 62.4 |
| | **EPI** | 719 | 260 | - | - | 449 | - | - | |
| | **COP** | 447 | 317 | - | - | - | - | - | |
| | **PAL** | 469 | 351 | - | - | 18 | - | - | |
| | **BER** | 2225 | 1563 | - | - | 162 | - | - | |
